# Using machine learning to overcome mosquito collections missing data for malaria modeling

**DOI:** 10.64898/2026.04.15.718796

**Authors:** Yasmin Rubio-Palis, Linjia Feng, Karena S. Liang, Chen Song, Songyuan Wang, Tymon Duchnicki, Xiao Zhang, Lelys Bravo de Guenni

## Abstract

Entomological surveillance plays a crucial role in areas where malaria remains endemic, yet gathering data on mosquito populations is often expensive and complicated, particularly in remote locations with challenging logistics and inconsistent sampling schedules. Access to extensive time series data on mosquito species at specific sites would greatly enhance insights into seasonal trends and the biting habits of vectors of malaria parasites. Gaps in mosquito count records pose a significant challenge for researchers and public health officials seeking to establish early warning systems and effective vector control programs. In this study, we apply quantitative machine learning techniques to address missing data in estimates of mosquito abundance collected from 2009 to 2016 in Bolívar State, Venezuela. We evaluated Linear Regression, Stochastic Linear Regression, K Nearest-Neighbor, and Gradient Boosting methods for imputing missing counts of *Anopheles* mosquitoes, employing a leave-one-out cross-validation strategy. Additionally, we developed a predictive malaria transmission model incorporating mosquito abundance and climate variables (El Niño 3.4 Index, rainfall, and mean air temperature) as covariates. Our generalized time series model forecasts malaria incidence of *Plasmodium vivax* and *Plasmodium falciparum* based on climate dynamics and imputed mosquito data. Model performance was assessed using root mean square error, mean absolute error, and mean absolute percentage error. The final results demonstrated that machine learning imputation significantly improved the accuracy and reliability of *P. vivax* malaria incidence predictions but failed to predict *P. falciparum* incidence. The study demonstrates that method choice significantly influences the reconstruction of seasonal abundance patterns and the performance of malaria incidence models. Nevertheless, the proposed models strengthen the foundation for targeted interventions and surveillance in endemic regions. Despite limitations in data continuity and coverage, the findings highlight the value of combining multiyear entomological data sets with robust imputation and sensitivity analyses to improve predictive modeling in resource-constrained, malaria-endemic settings.

## Introduction

Malaria stands as the most significant parasitic disease transmitted by mosquitoes. According to the World Health Organization [1], an estimate of 262 million malaria cases occurred across 80 endemic countries in 2024. Within the Americas, Venezuela reported an estimated incidence of 9.7-12.6 x 1,000 population at risk, significantly higher than the incidence reported by Brazil (3.08-3.43 x 1,000 population at risk), accounting for 17 percent of all malaria cases in the region. Notably, more than 70 percent of these cases originated from Bolívar State [1].

Mosquito control is a crucial intervention for preventing the transmission of malaria parasites. Nevertheless, effective surveillance and monitoring of mosquito vector populations are often constrained by the challenges of conducting regular collections across multiple locations, particularly in vast and remote regions. These logistical hurdles frequently result in incomplete data, leading to potentially inaccurate assumptions. Furthermore, efforts to model malaria transmission based on entomological data encounter significant limitations stemming from these data gaps.

A key mathematical approach to addressing this limitation involves leveraging machine learning techniques for imputing missing data. Numerous researchers have adopted such methods to estimate missing values effectively. For instance, Emmanuel et al. [2] provided a comprehensive review of the methodologies used for missing data imputation. Among the techniques employed, the K-nearest neighbor (KNN) method has demonstrated notable success. Their study also highlights that the performance of imputation methods, such as random forest-based algorithms and KNN, is highly dependent on the dataset being analyzed.

However, a potential limitation of the KNN method is its inability to consider the correlation between potential predictors, which can lead to unsatisfactory results [3]. To address such shortcomings, boosting algorithms have emerged as promising alternatives. Research by Cauthen et al. [4] demonstrated that boosting methods can outperform non-boosting approaches, especially in scenarios with high rates of missing values. Their study, which focused on imputing the number of dengue cases in India, highlighted the enhanced accuracy and robustness offered by boosting algorithms in handling complex, incomplete datasets.

The Extreme Gradient Boosting (XGBoosting) algorithm, an improved boosting method, has demonstrated superior performance in imputing medical data [3]. Additionally, both kernel-based and tree-based machine learning algorithms have been successfully applied to meteorological datasets [5]. In recent years, Recurrent Neural Network (RNN) models based on deep learning have become increasingly popular for imputing missing values in time series data, owing to their ability to extract and leverage temporal dependencies within datasets [6,7]. These studies typically rely on prediction error measurements, most commonly, root mean square error (RMSE) to evaluate algorithm performance. While machine learning methods are well-established for missing data imputation in various domains, their applicability to imputing mosquito abundance time series data remains largely unexplored. Expanding research in this direction could offer significant advances in handling incomplete entomological datasets, ultimately improving the accuracy of malaria modeling and the effectiveness of vector control strategies.

In this study, we systematically compare four distinct approaches for imputing missing data within incomplete time series of mosquito counts, focusing on the most abundant *Anopheles* species collected in a remote Amerindian community in southern Venezuela. Each imputation method leverages a range of potential climate drivers as predictors, including indices that capture the influence of the El Niño phenomenon. By applying these imputation strategies, we generate complete mosquito abundance time series, subsequently evaluating their performance using cross-validation error metrics. For both the most prevalent mosquito species (*Anopheles darlingi*) and for the aggregated dataset encompassing all species, we select the imputed time series that yield the lowest cross-validation error. *Anopheles darlingi* is the most efficient vector of malaria parasites in the neotropics [8,9,10], and the confirmed vector in the study area [11]; all the *Anopheles* species collected are potential vectors [10,12,13,14], therefore were included in the analysis. These optimally imputed mosquito abundance datasets are incorporated as potential predictors in our time series models of malaria incidence, with the aim of enhancing the accuracy of malaria transmission modeling in this challenging and data-limited context.

Beyond the challenges of imputing mosquito abundance data, numerous studies have explored the impact of climate drivers on malaria incidence across diverse regions. The importance of climate as a driving force of malaria transmission has been known for over a century [15]. The development and survival of *Anopheles* mosquitoes and the *Plasmodium* parasites are highly dependent on temperature, rainfall and humidity [16,17]. On the other hand, large scale climatic drivers as El Niño/La Niña (ENSO) phenomena are key in modulating the inter-annual variability of the precipitation worldwide [18]. The relation between ENSO and malaria epidemics has been studied in South America [19,20,21], and particularly in Venezuela [22,23,24,25,26]. The integration of climate variables as potential covariates in malaria prediction models has garnered increasing attention. For instance, Kifle et al. [27] demonstrated that rainfall data served as a reasonably good predictor of malaria incidence in Eritrea, highlighting the role of precipitation in shaping transmission dynamics. Similarly, Kumar et al. [28] introduced a Bayesian regression model incorporating climate predictors to successfully model malaria incidence in India. These findings underscore the potential of climate variables -such as rainfall, temperature, and humidity- as essential components in forecasting malaria risk and improving the precision of epidemiological models tailored to specific local contexts.

Importantly, mosquito abundance time series constitute a critical input for elucidating the causal pathways that drive fluctuations in malaria incidence. Previous works, such as Laneri et al. [29] have largely defined the causal landscape of malaria transmission in terms of climate variables, focusing on factors such as rainfall, temperature, and humidity. While this approach has yielded valuable insights, it may overlook the direct role that vector population dynamics play in modulating malaria risk.

The present study describes the missing-data imputation methods for *Anopheles* abundance time series, and the generalized time series models for the *Plasmodium vivax* and *P*. *falciparum* incidence in a malaria endemic region in southern Venezuela. We present the application of the different imputation approaches and compare their performance using a leave-one-out cross-validation (LOOCV) approach, together with an assessment of the generalized time series models for *P. vivax* and *P. falciparum* malaria.

After applying a variable-selection strategy based on predictive performance in a training set, the generalized time series models can be used to forecast future malaria cases in a testing set. These predictive models enhance our understanding of the interactions between climate, vector abundance and malaria incidence in remote areas.

## Materials and Methods

### Data Sources

#### Mosquito collection methods

Entomological surveillance in collaboration with local leaders from remote Amerindian communities was implemented in 2009 [30]. This action was demanded by the chief and members of the Ye’kwana community of Boca de Nichare (lat 06^∘^34.44’N, long 64^∘^49.39’W, altitude 59 msnm), Sucre Municipality, Bolívar State. In this area several studies have been conducted since 2005 with local leaders, and the demographic, ecologic, epidemiologic, hydrologic and geomorphologic characteristics have been described [11,30,31,32,33,34]. Local leaders were trained and supervised by one of the authors (YRP) in mosquito collections using the Mosquito Magnet Liberty Plus Trap (MMLPT) (American Biophysics Corporation, North Kingstown, RI). The training topics included mosquito identification, data collection, reporting, preservation, and shipping of specimens collected to the central laboratory in Maracay (Aragua State, Venezuela), where identification to species level and numbers collected were confirmed. The MMLPT device is battery operated and utilizes Carbon Dioxide (CO_2_), one of the products of the catalysis of propane gas, as an attractant together with Octenol. This trap has been previously evaluated in this region of southern Venezuela, and its efficiency to collect anophelines compared to human landing catches was determined [35].

#### Mosquito Data

Mosquitoes were collected monthly between 2009 and 2016, for 4 to 10 nights per month around the full moon from 18:00 to 06:00 hours. To estimate the biting pattern of the most abundant species, the trap was checked and the net with trapped mosquitoes was removed every four hours: 18:00 to 22:00 hrs; 22:00 to 02:00 hrs, 02:00 to 06:00 hrs. The total number of individuals was recorded, and mosquito species were identified. The most predominant species were *Anopheles darlingi* (darl.1), *Anopheles oswaldoi* (osw.5), *Anopheles goeldii* (goel.6), and other species with a less important presence in the area included *An. braziliensis, An. triannulatus, An. benarrochi* and *An. mattogrossensis*. A time series of the estimated mean number of mosquitoes by combining all *Anopheles* species collected was also considered in the analysis and will be referred to as *All* in the text. The main assumption is that *Anopheles* abundance and species diversity is representative of the municipality.

#### Mosquito Abundance Time Series

The monthly mosquito abundance time series for the most abundant species (*An. darlingi, An. goeldii,* and *An. oswaldoi*), and the monthly time series combining all *Anopheles* species collected (*All*) is presented in Fig 1.

**Fig 1.**
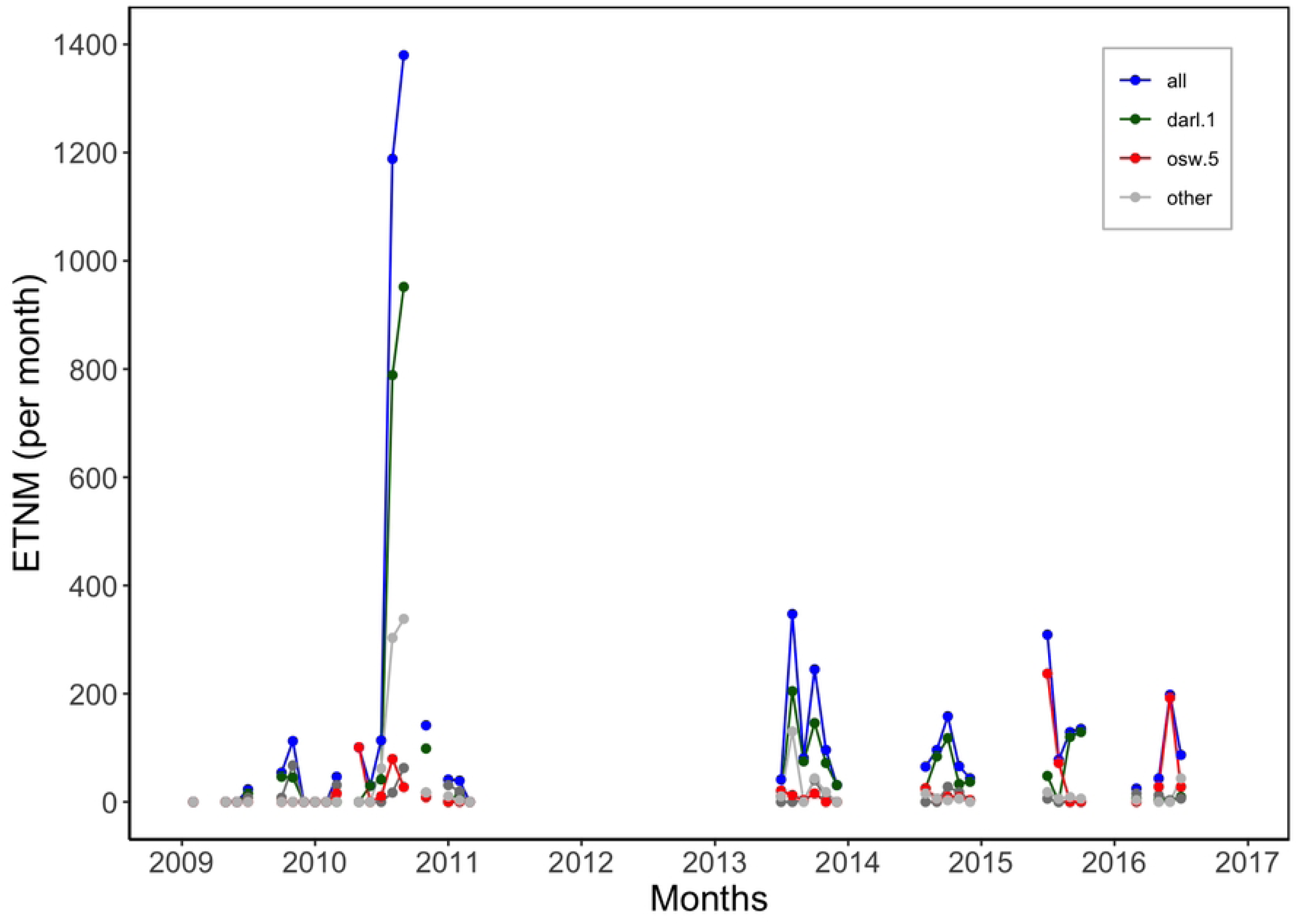
Estimated total number of mosquitoes (ETNM) collected per month. All = total number of mosquitoes collected of all *Anopheles* species; darl.1 = *Anopheles darlingi*; goel.6 = *Anopheles goeldii*; osw.5 = *Anopheles oswaldoi*; Other = total number of other *Anopheles* species collected.

All *Anopheles* species identified during the surveys are considered potential vectors [9,10,11,36] and, hence were included in the time series and other analysis. These values were calculated by adding the total number of mosquitoes collected per night during three consecutive four-hour periods from 18:00 to 06:00 hours. The average number of mosquitoes per night was calculated by adding the total number of mosquitoes for all nights of observation and dividing by the total number of nights in a particular month. This estimate was converted into a monthly estimate by multiplying the mean number of mosquitoes per night by *ndays*, where *ndays* is the total number of days in each month.

A distinctive feature of this data set is the important proportion of missing data. The proportion of missing observations (or months without collections) is 60.4% of the total number of observations. As shown in Fig 1, there are important data gaps, especially during the years 2012-2013 due to travel limitations to the collection site, including economic restrictions and the shortage of fuel.

#### Mosquito Abundance and Biting Activity

Mosquito abundance and biting activity can vary from year to year, depending on the climatic drivers. It can vary between species and could also vary between different collection periods (night hours) [11,37,38,39]. To determine the significance of these factors on the mosquito biting behavior, a three-way ANOVA model was fitted to the mean number of mosquitoes by year, species and collection time, including also the interactions among all the three different factors. In this case, a normality Shapiro-Wilks test was not rejected. The variable representing collection times (with three intervals: 18:00 to 22:00 hrs; 22:00 to 02:00 hrs, 02:00 to 06:00 hrs) and its interaction with the other two factors, resulted as non-significant, indicating that there are not significant differences in the number of mosquitoes among these distinct mosquito collection periods. For this reason, this factor was not accounted for in the analysis herein.

#### Malaria Data Cases

Monthly malaria cases due to *P. vivax* (PV) and *P. falciparum* (PF) were obtained from the Malaria Program report for the Sucre Municipality. Cases resulting from mixed infections associated with both parasite species were also available and added to the corresponding number of malaria cases for each parasite. Data on malaria cases for the study location (Boca de Nichare) was not available.

Population data for the Sucre municipality was available from 2009 to 2017 (Instituto de Salud Pública, Bolívar State). The monthly *P. vivax* and *P. falciparum* incidence were estimated by dividing the total number of cases of each parasite species by the population per 1,000 population. The variables *P. falciparum* incidence (PF), and *P. vivax* incidence (PV) were used as response variables in the malaria model described in the methodology section. Time series of this data set are shown in Fig 2.

**Fig 2.**
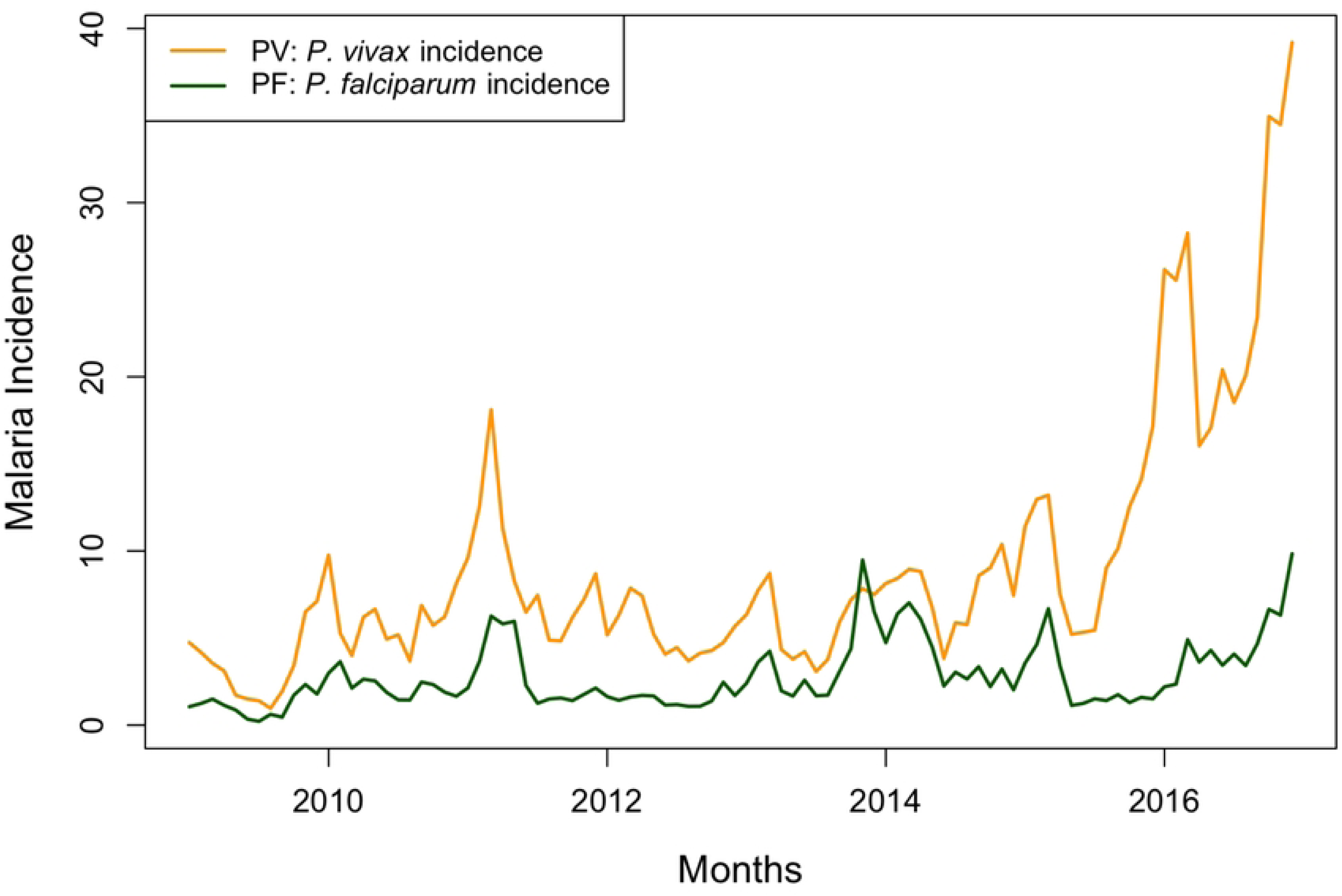
*Plasmodium vivax* and *Plasmodium falciparum* incidence, Sucre Municipality, Bolívar State, Venezuela, 2009-2017.

#### Climate data

The climate of the study region has been classified as a Tropical Savanna Climate (Aw) according to Köppen-Geiger climate classification system, with annual average temperatures of approximately 23.46°C and average annual precipitation of 2,764 mm/yr.

Nearby meteorological stations with long term time series data are unavailable for the study site. Historical climate information is readily available from open data sources, including large scale oceanic-atmospheric indices depicting the temporal variability of the El Niño-Southern Oscillation (ENSO) index.

Rainfall and mean air temperature data were downloaded from the World Bank data portal https://climateknowledgeportal.worldbank.org/. Fig 3 shows the monthly historical time series for the study region.

**Fig 3.**
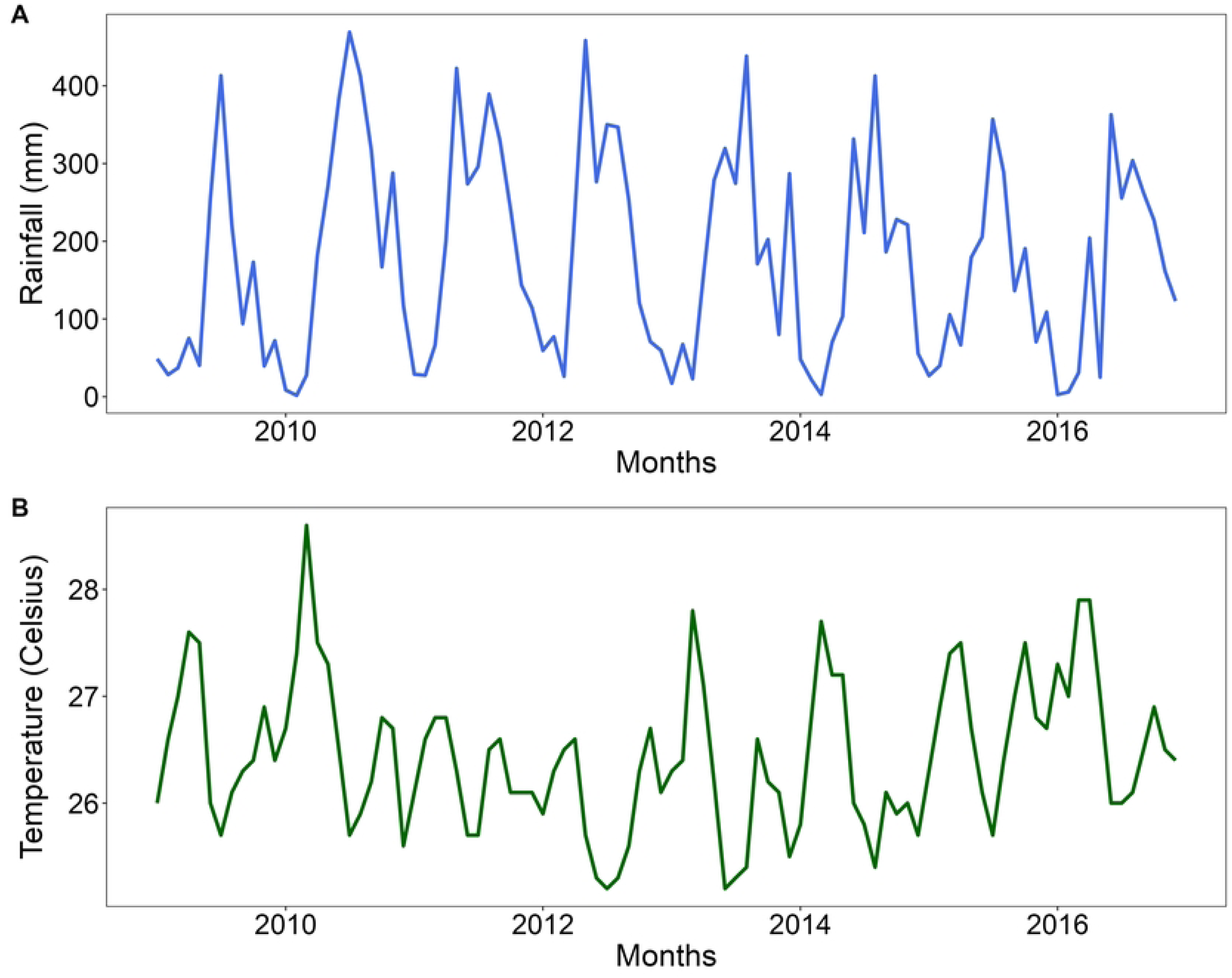
Rainfall and Mean Air temperature time series during the period 2009-2016. A. Total Rainfall in mm. B. Mean Air Temperature in degrees Celsius. Boca de Nichare, Bolívar State, Venezuela.

The El Niño 3.4 Index (Nino3.4) was downloaded from the National Oceanic and Atmospheric Administration (NOAA) website https://psl.noaa.gov/enso/data.html. This index depicts the values of the Sea Surface Temperature (SST) Anomalies for the Pacific Ocean region with extents (5N-5S, 120W-170W) (S1 Fig).

Fig 4 shows the estimated total number of mosquitoes (ETNM) per month time series together with the anomalized rainfall and the anomalized temperature.

**Fig 4.**
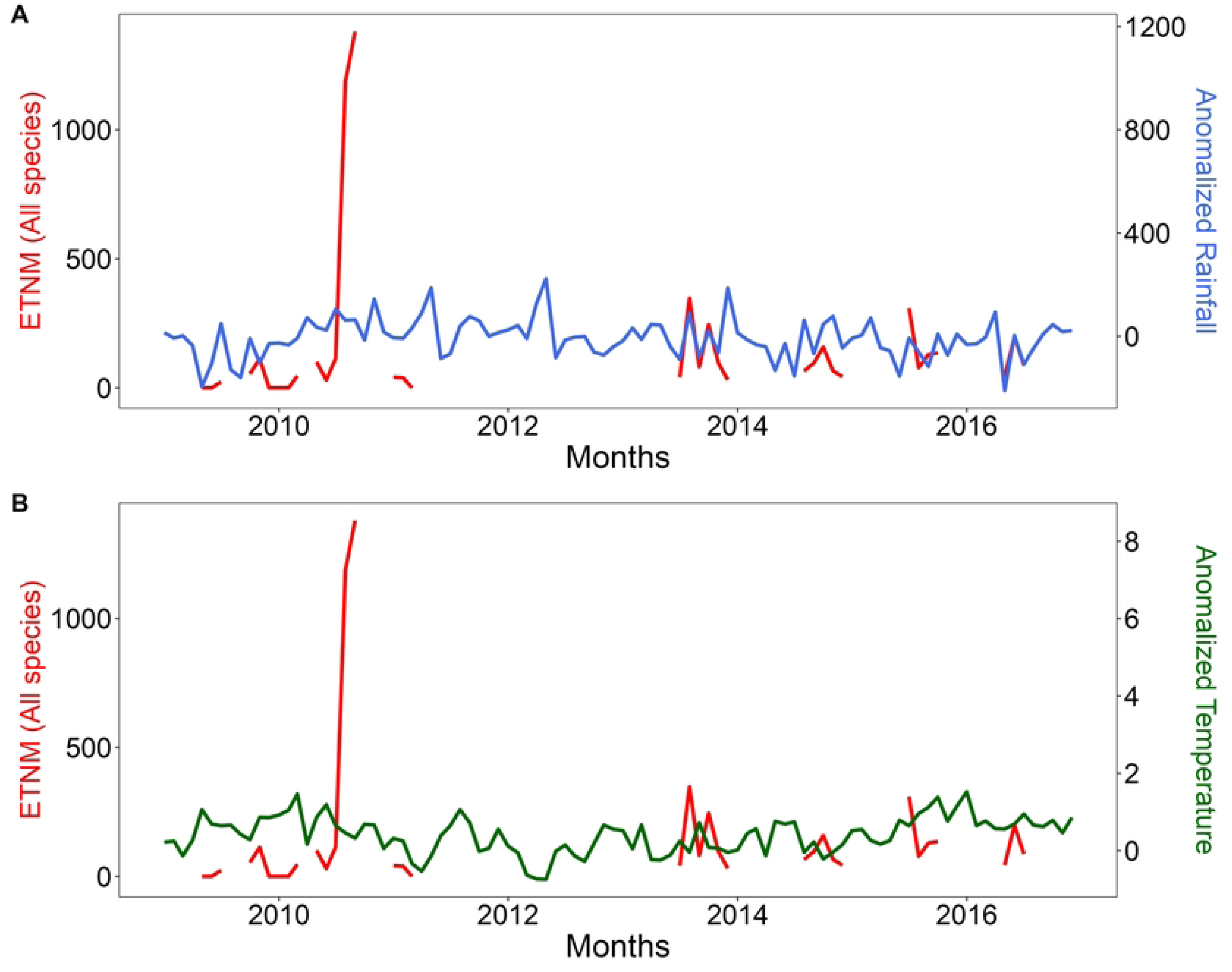
Estimated total number of mosquitoes (ETNM) per month for All *Anopheles* species (red line) compared with climate data. A. Anomalized mean monthly rainfall in mm (blue line). B. Anomalized mean monthly air temperature in degrees Celsius (green line). Boca de Nichare, Bolívar State, Venezuela (2009-2016).

Anomalized rainfall and temperature time series were calculated by subtracting the long-term monthly means from each monthly value. The anomalization procedure filters out the series’ seasonal components while the monthly and inter-annual variability remain as dominant components. The long-term monthly means were calculated using data from the period 1981 to 2010.

To compare the estimated mean number of mosquito time series with the ENSO data, Fig 5 overlaps the different ENSO occurrence modes with the mosquito data before imputation. The highlighted peak in the mosquito times series occurs during a weak La Niña phase. These maxima occur during a positive rainfall anomaly period, also preceded by a positive air temperature anomaly.

**Fig 5.**
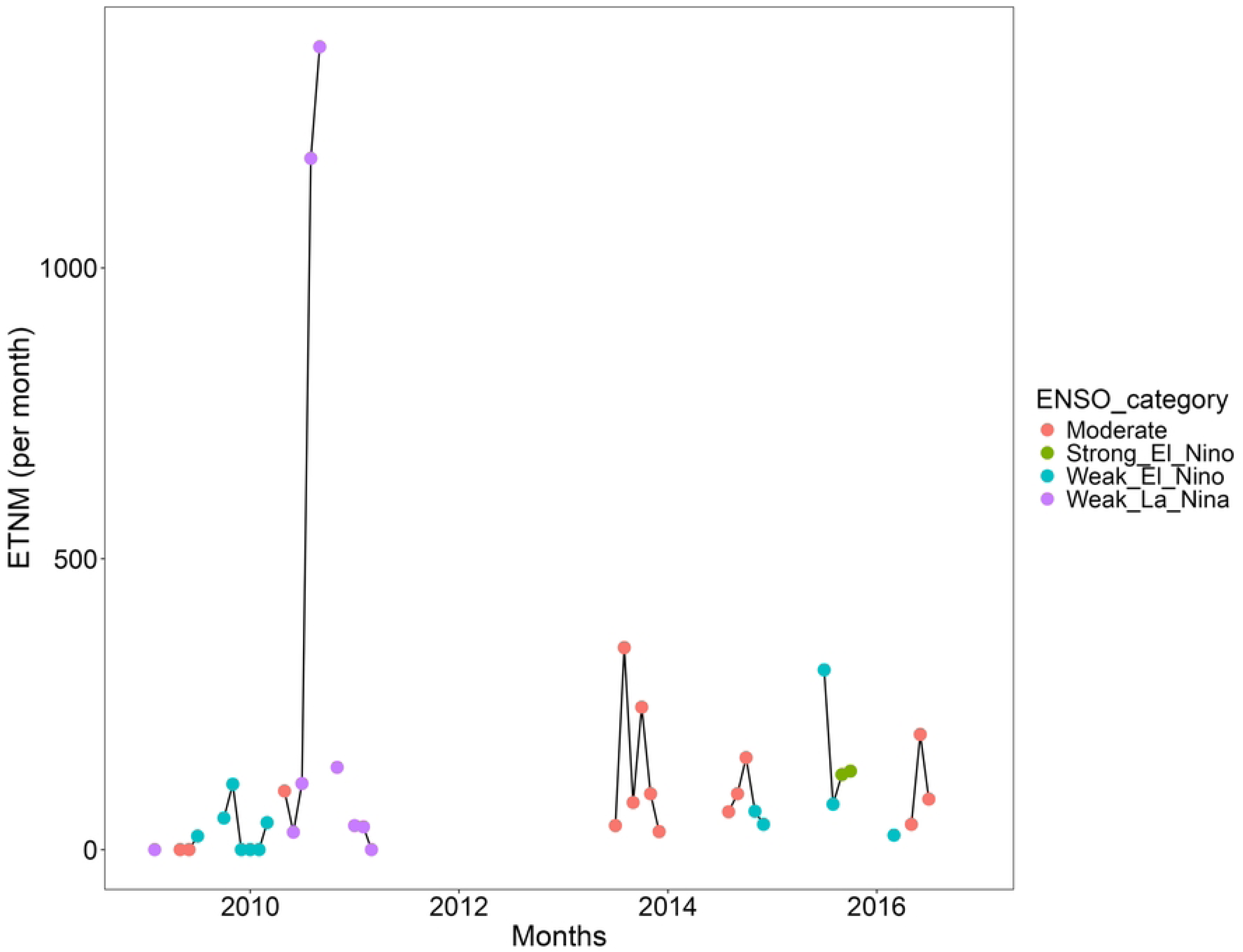
Estimated Total Number of Mosquitoes (ETNM) per Month and El Niño.

### 3.4 index for the period 2009 – 2016. Boca de Nichare, Bolívar State, Venezuela

A summary of the climatic variables during the study period 2009-2016 is presented in Table 1.

**Table 1.**
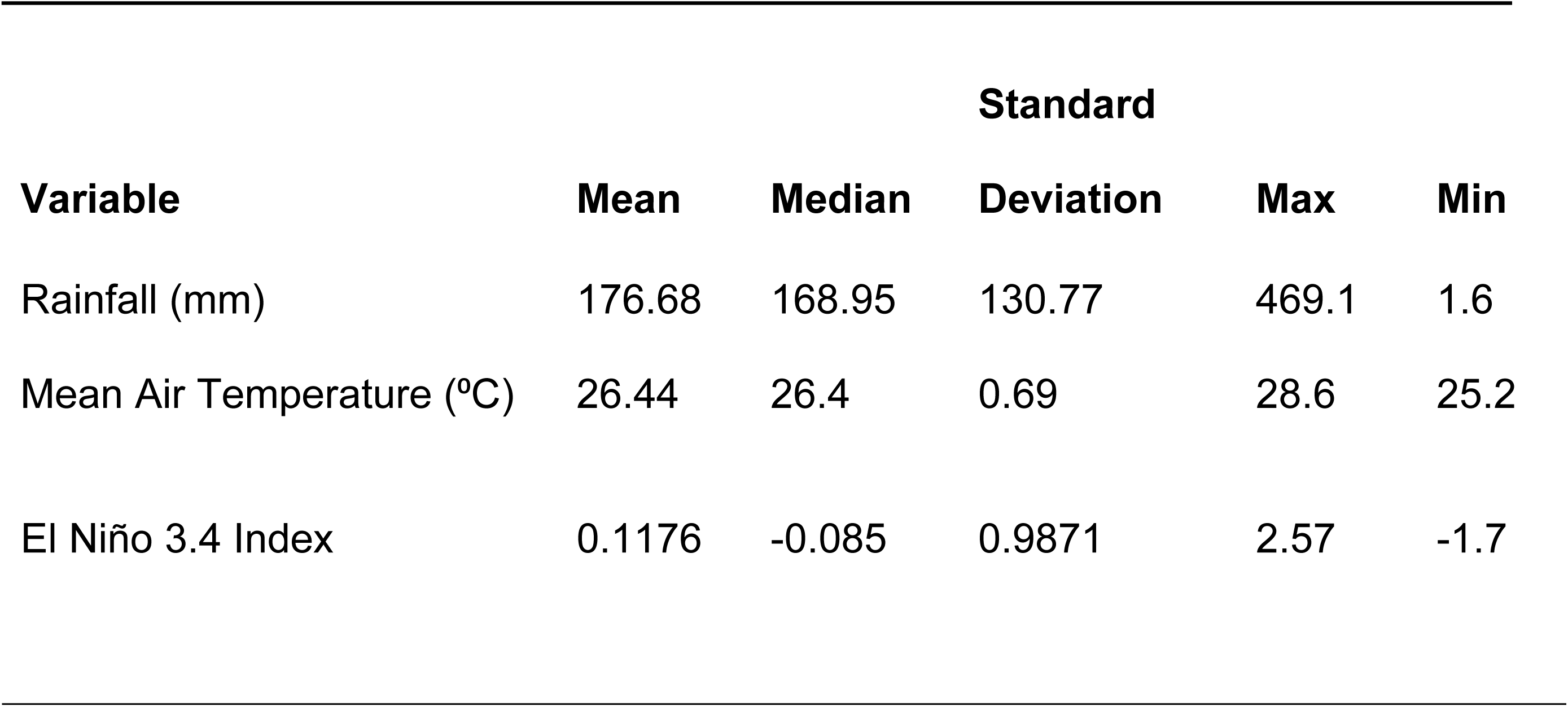
Summary monthly statistics for climatic variables Rainfall, Mean Air Temperature and El Niño 3.4 index during years 2009-2016.

### Methodology

In this section, we describe the different methods used for imputation of mosquitoes (*An. darlingi* and all *Anopheles* species) time series missing data and present the generalized time series model used to model malaria incidence.

#### Missing Data Imputation Methods

Missing data handling methods are usually adapted to the nature of missing data mechanisms. The missing data mechanisms are related to the dependence or not of the missing data on the observed values. Emmanuel et al. [2] and Sterne et al. [40] propose classifying missing data according to their type: missing completely at random, missing at random and missing not at random. The type of missing data we are dealing with in this study can be classified as missing completely at random because missing values do not depend on the observed or unobserved data, but rather on the ability to collect information in a timely manner if economic and/or logistic resources (gasoline, propane gas, trap functioning) are readily available.

Deleting missing data is one method of handling missing values called the Complete Case Analysis. This procedure is rarely applied because deleting missing data points (Listwise deletion) can significantly reduce our sample size for model training purposes, and therefore, accuracy would be a concern.

Using imputation methods as regression imputation allows for estimating missing values by using other variables as covariates or predictors. In this research, different approaches were used for missing data imputation. The following methods will be discussed and compared:

Linear Regression (LR), Stochastic Linear Regression (SLR), K Nearest-Neighbor (KNN) and Gradient Boosting (GB).

The imputation methods LR, SLR, KNN and GB were implemented in Python (Google Colab tool) using packages fancyimpute, sklearn and xgboost.

A brief description of each method and the steps for their implementation are described below.

### Linear Regression and Stochastic Linear Regression Imputation

Regression imputation is a technique used to replace missing values *Y*_*m*_in a data set based on a set of predictors or attributes *X*_1_,*X*_2_,…,*X*_*p*_. It can be classified into two types: deterministic regression imputation and stochastic regression imputation. Deterministic regression imputation predicts missing values using the estimated regression model:

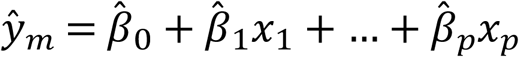

where *β̂*_*i*_ are the parameters of the estimated regression model using least squares, and *ŷ*_*m*_ are the estimated mean values of the response variable *Y* given a set of values for the predictors *x*_1_,*x*_2_,…,*x*_*p*_. Estimation of the parameters *β*_0_,*β*_1_,…,*β*_*p*_ is based on the observed values *Y*_*o*_ and the corresponding observations for the predictors (complete cases). However, this method uses a point estimate that does not consider any random variation (i.e., an error term) around the regression model, leading to imputed values that may be too precise resulting in an overestimation of the correlation between *X*_*i*_ and *Y*.

To address this limitation, stochastic regression imputation was used. This method incorporates a random error term into the predicted value:

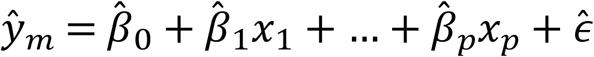

where *ε̂* is a simulated random error term from a normal distribution with zero mean and variance estimated using least squares. By incorporating this error term, stochastic regression imputation can reproduce the correlation between *X*_*i*_ and *Y* more accurately.

Using regression imputation, we used the Leave-One-Out Cross-Validation (LOOCV) method to calculate the average RMSE for the evaluation of imputation performance. LOOCV is a special type of k-fold cross-validation in which the number of folds *k* is equal to the number of observations *N* in the data set. Under this approach, each observation will be treated as a validation set once, and the rest (N-1) observations will be used as a training set. This method aims at reducing bias and preventing over-fitting, and it is useful for data sets with a small number of observations.

#### K-Nearest Neighbor method

This algorithm falls under the umbrella of supervised learning methods. It can be used for both regression and classification problems. It is considered a *Lazy learner* algorithm [41] and does not assume a probability distribution of the data. Instead, KNN assumes that similar inputs have similar outputs, and it will impute the missing data with the average value amongst its *k* most similar inputs. KNN depends heavily on the parameter *k*, which is the number of nearest neighbors to be used for imputation. The range of *k* values used in our imputation process varied from 1 to 40.

The KNN algorithm uses a distance measure to detect the closest data points to a target vector *x*_0_in a neighborhood, for which a missing characteristic *y*_0_should be imputed. It is also important to note that the *K* neighbors do not have any missing features. Examples of distance measures are the Minkowski distance, Manhattan Distance, Cosine Distance, Jaccard Distance, Hamming Distance and Euclidean distance [2]. Among these distance measures, Euclidean distance is commonly used in practice. The equation is:

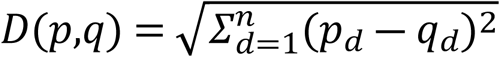

where *D*(*p*,*q*) is the Euclidean distance, *p* and *q* are two data points in an n-dimensional space and *n* is the dimension of the space.

The imputed value at *x*_0_ (*ŷ*_0_) is the average of all responses *y*_*i*_ in the neighborhood:

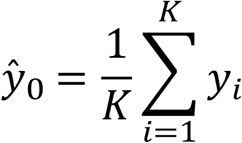

The LOOCV method (Leave-One-Out Cross-Validation) is used to select the optimum value of *K* by comparing the value of the root mean square error (RMSE) for each number of neighbors *K* and selecting *K* that minimizes RMSE.

#### Gradient Boosting Method

The Gradient Boosting method (GBoosting) is also a supervised learning algorithm where a training data set is used to predict a response *y*_0_ from an input vector of features *x*_0_. It can be used both in a regression context and as a classifier. The final prediction is an ensemble of decision trees constructed in an iterative way. Each tree is considered a *weak learner,* and the algorithm uses the steepest descent method to move into the direction of minimizing a loss function at each step. The Gradient Boosting method employs regularization techniques to prevent overfitting, and it can handle missing data automatically. The Gradient Boosting model is flexible for data with a large missing data rate and small size, and it is also useful for minimizing bias error.

A wide range of hyperparameters must be specified for the Gradient Boosting model. The hyperparameters used in the imputation process are max_depth (maximum depth of a tree), min_samples_leaf (minimum number of observations in a leaf), and learning_rate (how quickly the model learns). The LOOCV method is used to select the best combination of the three parameters with the lowest RMSE.

The gradient boost algorithm objective is to find an approximation of the function F(x) that can minimize the expected value of the loss function L(y, F(x)) [42]:

In the first step, we want to minimize the loss function *L*(*y*_*i*_,*α*), which represents the difference between the actual values and the predicted values. A constant prediction of the model *F*_0_(*x*) is given by:

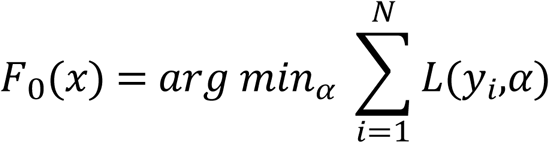

Where *α* is the value that minimizes the loss function 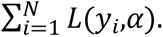

In subsequent steps (*m* = 1,…,*M*) the following expression is minimized:

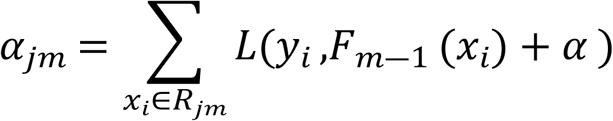

At each step, several regression trees are created on the residuals of the model. These residuals are calculated as follows:

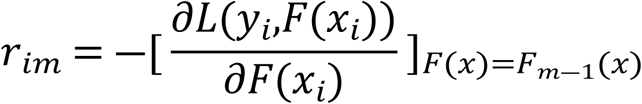

where *i* = 1,…,*N*; and m is the index of each tree.

The derivative is taken with respect to the previous prediction *F*_*m*―1_, and the result is proportional to the residual value at the step *m* ― 1.

In step *m* the prediction *F*_*m*_(*x*) is updated as:

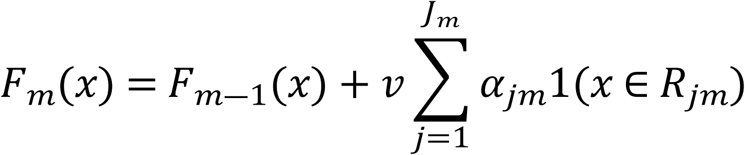

*j* is the index of each terminal node (leaf); 1(.) is the indicator function and *J*_*m*_ is the total number of leaves in the tree. *R*_*jm*_ represents each terminal node and *v* ∈ (0,1) is the learning rate.

### Generalized Time Series Model for Malaria Incidence

One method for modeling malaria cases with covariates is using a generalized linear model for count time series. Let {*Y*_*t*_:*t* ∈ *N*} be a count time series and {*X*_*t*_:*t* ∈ *N*} be a time-varying *r* dimensional covariate vector. We then model the conditional mean *E*(*Y*_*t*_|*F*_*t*―1_) of the time series by a process {*λ*_*t*_:*t* ∈ *N*} such that *E*(*Y*_*t*_|*F*_*t*―1_) = *λ*_*t*_, where *F*_*t*_ is the history of the joint process {*Y*_*t*_,*λ*_*t*_,*X*_*t*+1_:*t* ∈ *N*} up to time *t* and includes the information of covariates at time *t* + 1. Furthermore, let *g*:*R*^+^→*R* be a link function and *g̃*:*N*_0_→*R* be a transformation function. The count time series model, as proposed in Liboschik et al. [43], is of the form:

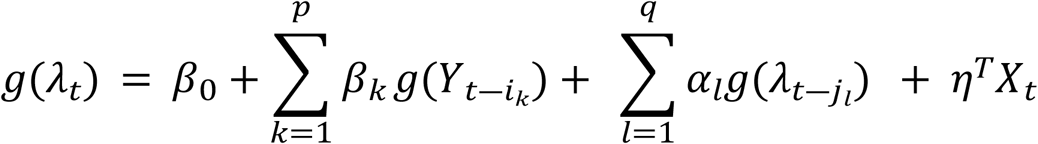

Here *η* is the parameter vector of the covariate effects, and *β*_0_ is the intercept. We can regress on past observations of the time series by defining a set *P* = {*i*_1_,*i*_2_,…,*i*_*p*_} of integers where 0 < *i*_1_ < *i*_2_… < *i*_*p*_ < ∞ and then regressing on lagged observations 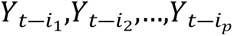 with effects *β*. We are also able to regress on lagged conditional means by defining a similar set *Q* = {*j*_1_,*j*_2_,…,*j*_*q*_} and regressing on conditional means 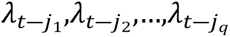 with effects *α*.

The model is fitted using quasi-conditional maximum likelihood estimation. If the link function is the identity function, then *g*(*x*) = *g̃*(*x*) = *x*. Otherwise, if the link function is logarithmic, then *g*(*x*) = *log*(*x*) and *g̃*(*x*) = *log*(*x* + 1). Furthermore, it must be assumed that *Y*_*t*_|*F*_*t*―1_ ∼ *Poisson*(*λ*_*t*_) or *Y*_*t*_|*F*_*t*―1_ ∼ *Negative Binomial*(*λ*_*t*_,*ϕ*) where *ϕ* ∈ (0,∞) is an additional dispersion parameter. For details on parameter estimation, refer to the documentation of the R package *tscount* [43].

## Results

### Climate data and mosquito time series dependence

Sample cross-correlation functions were calculated to understand the association between the estimated total number of mosquitoes per month and the climatic variables. As mentioned, the rainfall and temperature data time series were anomalized to partially remove the strong seasonal pattern from these climate variables.

Fig 6 shows the sample cross-correlation function between the estimated total number of mosquitoes (ETNM) per month and the monthly values of the El Niño index (Niño3.4), the anomalized mean temperature, and mean monthly rainfall for all *Anopheles* species.

**Fig 6.**
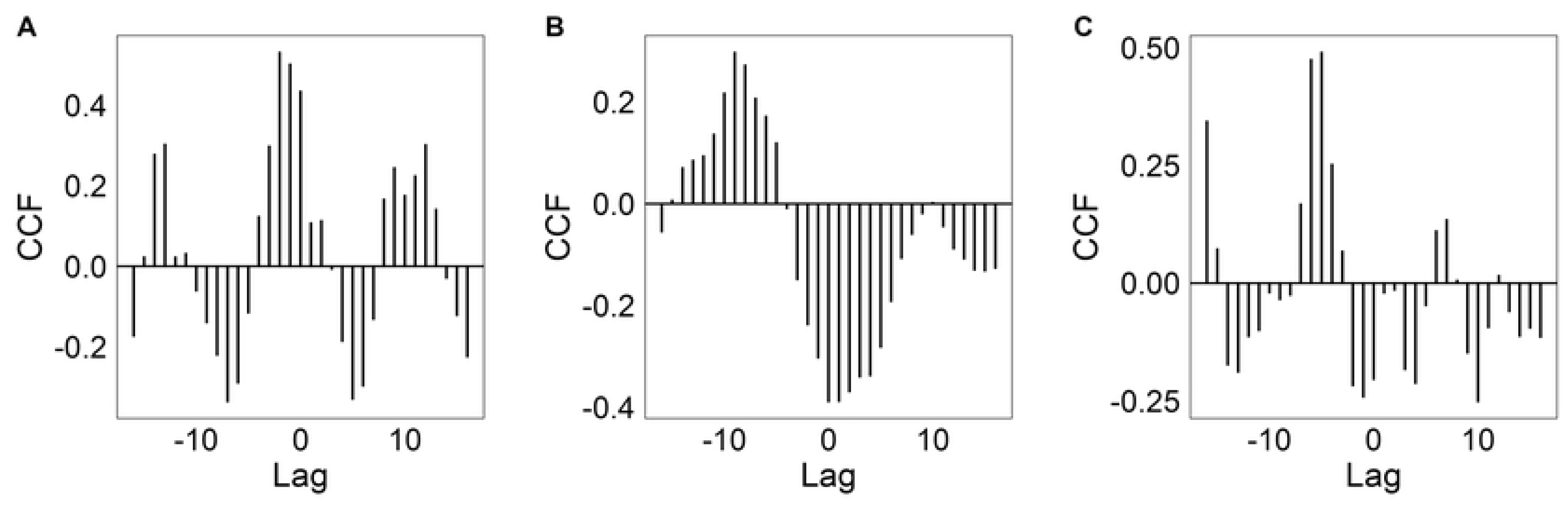
Cross-correlation between the estimated total number of mosquitoes (ETNM) for all *Anopheles* species time series and the Niño3.4 index (A), the monthly mean temperature (B), and monthly rainfall (C).

By inspecting the negative lag portion of the figures, we can detect the leading lag for which the mosquito data is significantly correlated with the climate variables.

The maximum correlation in absolute value within a year occurs for lags of 9, 6, and 2 months respectively. There is a significant positive association between mosquito abundance and El Niño index, temperature, and rainfall after 9, 6, and 2 months, respectively.

A similar calculation was repeated for the most abundant species, *An. darlingi* (darl.1) and the same lagged significant effect of climate drivers on mosquito abundance were observed.

### Imputation methods comparison

#### Linear regression and stochastic linear regression

The linear regression model was fitted by least-squares, using all non-missing data values (referred to as *complete cases*) as the training data set. This fitted model was subsequently used to impute missing values. The model was based on predictors including temperature and rainfall anomalies and the El Niño3.4 index. These predictors were initially considered without any lagged effects (i.e., no time delays). After analyzing the cross-correlations (Fig 6), lagged versions of the climate predictors were incorporated into the model, using the selected lags exhibiting the highest cross-correlation with the estimated total number of mosquitoes (ETNM) response variable. We then made a comparative analysis of model performance under these two scenarios: using lagged predictors and using non-lagged predictors.

Our primary focus was on the imputation of data associated with all mosquito species, obtained by aggregating mosquito counts from all species. In addition, we examine data imputation for each species separately. For assessing the prediction error after imputation of missing values, the following approach was employed:

1. Due to the limited availability of non-missing data, a simple random sampling of the observed data with replacement was implemented. The number of samples from this sampling process was equivalent to the number of missing data values. Multiple training data sets were initially generated by repeatedly applying the random sampling method for *K* iterations.
2. A linear regression model was fitted to each training sample using least-squares. To assess model performance, a leave-one-out cross-validation (LOOCV) error was calculated for each of these training data sets, and the average root mean square error (RMSE) was obtained.
3. In our analysis, we observed that using lagged climatic variables as predictors resulted in a better model performance (RMSE= 141.40) compared to using climatic variables without lagging (RMSE=142.69) (S2 Fig).
4. The same procedure was followed for the stochastic model: climatic variables as predictors were considered before lagging (no lag) and after lagging, and model performance was compared in these two cases. The stochastic regression approach used the same random sampling function as the linear regression model to impute missing values. It generated random predictions based on the regression model, considering the prediction standard error to fill in the missing values. Random predictions were generated from a normal distribution centered at the estimated mean predicted value, with a standard error equal to the standard deviation of the residuals for each estimate. If the random prediction was greater than zero, it was used as the imputed value for that specific missing value. To evaluate the performance of the stochastic imputation method, we used the leave-one-out cross-validation (LOOCV) method, resulting in a list of RMSE values for each training sample that were averaged for all iterations. Our analysis indicated that utilizing lagged climatic variables mean (RMSE= 0.13) led to a slightly better performance compared to using climatic variables without lagging mean (RMSE=0.10) (S3 Fig).

#### K-Nearest Neighbor (KNN)

The model was trained using the leave-one-out cross-validation method (LOOCV) to adjust the parameter *K* (number of nearest-neighbors). Climatic variables as predictors were considered before lagging (no lag) and after lagging, and the model performance was compared in these two cases. The following steps were followed:

The LOOCV method was used to select the best value of the *K* parameter. A KNN model was fitted to all five species for values of *K* in the range of 1 to 40. The value of *K* with the lowest RMSE for the LOOCV method was *K* = 16 in *An. darlingi* (darl.1). Fig 7 shows a comparison of the RMSE for each value of *K* for *An. darlingi* (darl.1) and All mosquito species.

**Fig 7.**
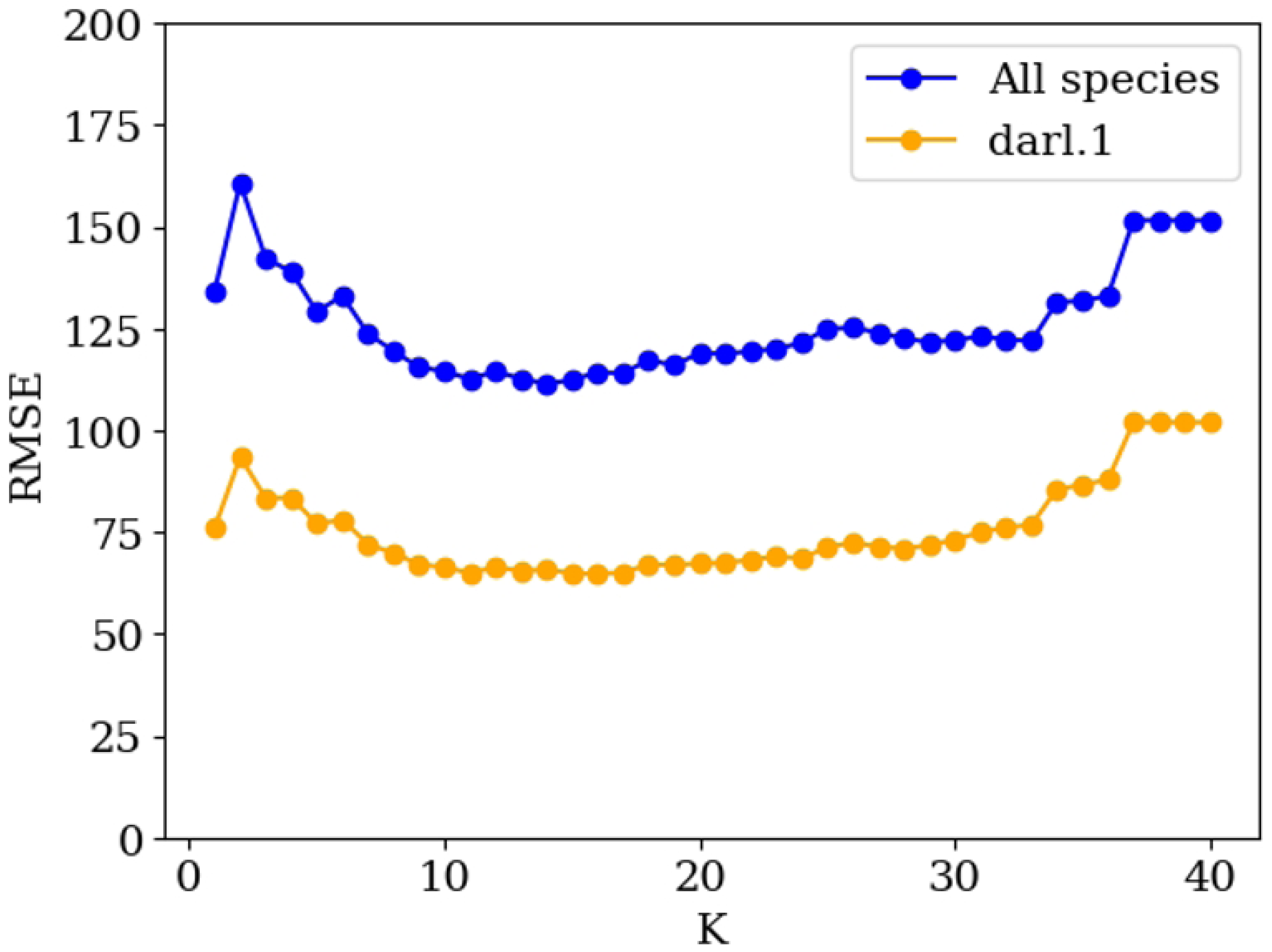
Comparison of the leave-one-out cross-validation (LOOCV) root mean square error (RMSE) for each *K* value for *Anopheles darlingi* (darl.1) and for All *Anopheles* mosquito species.

The KNN imputation model was constructed for *An. darlingi* (darl.1) and for All species with the number of neighbors *nneighbors*=16 and a data *train*:*test* split ratio of 8:2. The model was run 1,000 times, and the root mean square error (RMSE) was calculated for each iteration. A better performance of the methodology was found when using lagged climatic variables (RMSE= 126.83) in comparison to using climatic variables with no lagging (RMSE= 164.47) for darl.1. Fig 8 shows a comparison of the RMSE for each iteration using climatic variables before and after lagging. These results confirm better predictability of the fitted model when using lagged climate predictors.

**Fig 8.**
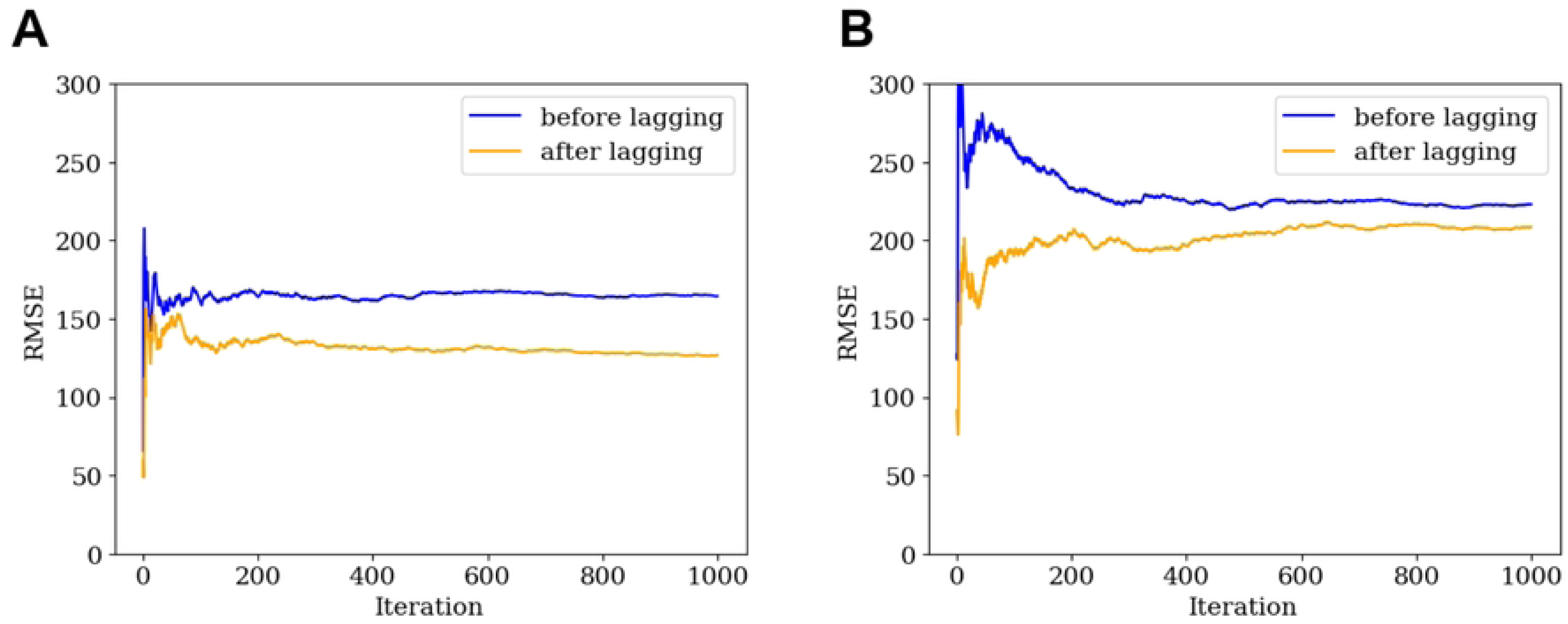
Comparison of root mean square error (RMSE) for K-nearest neighbor (KNN) method in each run, using climatic variables before and after lagging. A: *Anopheles darlingi* (darl.1); B: All *Anopheles* species collected.

#### Gradient Boosting (GB)

The model was trained using the leave-one-out cross-validation (LOOCV) method by tuning three parameters in the Gradient Boosting algorithm, which include *max depth*, *min samples leaf*, and *learning rate*. Climatic variables as predictors were considered before lagging (no lags) and after lagging in the same way as with previous imputation methods, and model performance was compared in these two cases. The following steps were followed:

1. The LOOCV method was used to select the best value of the parameters mentioned above. The GB model was fitted to species *An. darlingi* (darl.1) and All species for different values of the hyperparameters. We used a grid search to tune the hyperparameters, aiming to find the optimal combination of three hyperparameters. The hyperparameter values with the lowest RMSE for LOOCV were *min samples leaf* = 1, *learning rate* = 0.1, *max depth* = 10.
2. The Gradient Boosting imputation model with the default hyperparameters of GB was constructed for *An. darlingi* (darl.1) to compare the performance between lagged and no-lagged climate variables. The dataset was split into training and testing sets at an 8:2 ratio. The model was run 100 times and the root mean square error (RMSE) was calculated for each iteration. The methodology performed better when using lagged climatic variables (RMSE= 163.75) compared to non-lagged climatic variables (RMSE= 217.44) for darl.1. Fig 9 shows a comparison of the RMSE for each iteration using climatic variables before and after lagging. These results again highlight the advantage of using lagged predictors over non-lagged ones.

**Fig 9.**
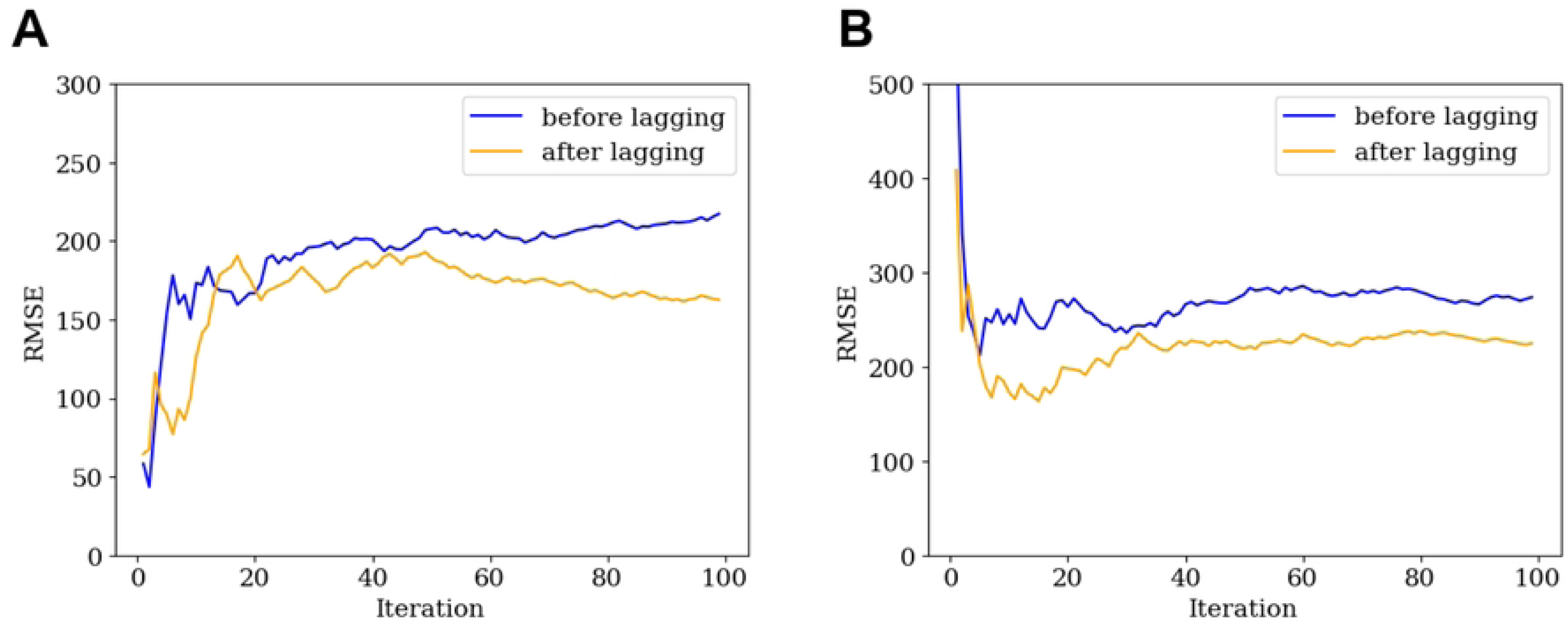
Comparison of root mean square error (RMSE) for the Gradient Boosting (GB) method in each run. Using climatic variables before and after lagging. A: *Anopheles darlingi* (darl.1); B: All *Anopheles* species collected.

Fig 10 show the contrast between the original data (in blue) and the imputed data (in yellow) for the four different imputation methods used for the species *An. darlingi* (darl.1).

**Fig 10.**
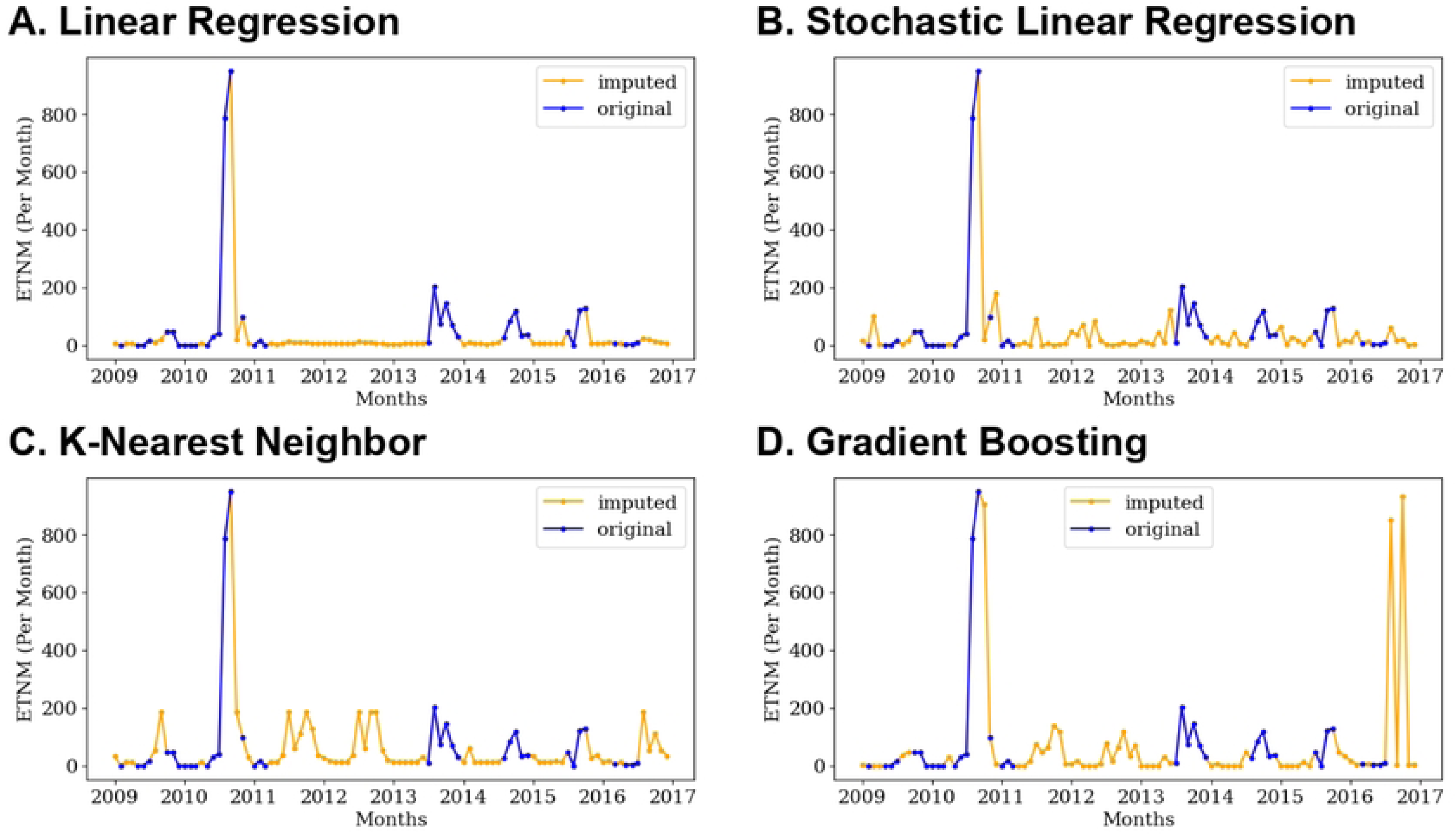
Estimated total number of *Anopheles darlingi* abundance (ETNM) time series imputation methods. A. Linear Regression (LR). B. Stochastic Linear Regression (SLR). C. K-Nearest Neighbor (KNN). D. Gradient Boosting (GB).

Important differences can be distinguished among the different methods, especially between linear regression imputation and the remaining methods. These observed differences are not surprising given the variety of imputation approaches used. A comparison of the prediction errors based on the non-missing or complete case data values can be assessed by estimating the RMSE using a LOOCV method for each imputation approach. These results are presented in Table 2. Table entries are relative numbers, calculated after dividing by the mean number of mosquitoes for each species.

**Table 2.** Leave-One-Out Cross Validation (LOOCV) method errors^a^ for the different imputation methods and mosquito species.

| *Anopheles* |  |  |  |  |
| --- | --- | --- | --- | --- |
| Species | Imputation Method |  |  |  |
|  | KNN | G. Boosting | Linear R. | Stochastic R. |
| All<sup>b</sup> | 0.7638104005 | 0.0000000297 | 0.968946389 | 0.000917769 |
| *An. darlingi* | 0.7727904964 | 0.0000000539 | 0.972812828 | 0.001148532 |
| *An. goeldii* | 0.8078696262 | 0.00000012 | 1.308744024 | 0.002081277 |
| *An. oswaldoi* | 1.345153286 | 0.000000627 | 1.503379349 | 0.001657576 |
| Other | 1.073280571 | 0.00004271 | 1.120539972 | 0.001491594 |

<sup>a</sup>The root mean square error (RMSE) values were calculated using the complete case observations (non-missing cases).
<sup>b</sup>All *Anopheles* species collected.
<sup>c</sup> Other than *An. darlingi*, *An. goeldii* and *An. oswaldoi* species collected.

The GB methodology shows the best predictive performance with the lowest testing errors in all cases. The same analysis was done for All *Anopheles* species, with similar results (S4 Fig).

### Malaria Incidence Time Series Model

#### Malaria Incidence Model Covariates

For the generalized linear time series count model described above, we consider several potential covariates for inclusion in the model: monthly rainfall, mean monthly temperature, ENSO index (El Niño 3.4), and an imputed mosquito count time series. Initially, we selected the imputed mosquito time series with the lowest LOOCV error which corresponds to the Gradient Boosting approach (Table 2).

Additionally, we considered lagged versions of each predictor. To select the optimal lag, we calculated the rank cross-correlation of the malaria incidence time series and each climatic anomalized predictor after removing the long-term monthly means. The selected lag was the one that resulted in the highest cross-correlation between the anomalized climatic time series and malaria incidence. For rainfall, a two-month lagged time series was considered based on the time lag that takes into account the time for creating habitats for the development of the aquatic stages of *Anopheles* mosquitoes (from eggs to adults), the buildup of mosquito populations, blood feeding interval and the development of the parasite inside the mosquito (sporogony) [37,44,45,46,47].

Figs 11, 12, 13, and 14 show the sample rank cross correlation functions of the *P. vivax* and *P. falciparum* incidence with rainfall, mean air temperature, ENSO index, and imputed All *Anopheles* species time series. The lag values for the negative portion of the graphs with the highest rank cross-correlations in absolute value are shown in Table 3. Note that cross-correlations between rainfall and *P. vivax* incidence are nonsignificant at any lag. However, the negative lag corresponding to the highest cross-correlation (in absolute value) was selected as a model covariate. This lag varied between 4 and 5 months for rainfall, 8 months for temperature, 16 and 17 months for the ENSO index, and 4 to 6 months for mosquito counts (Table 3).

**Fig 11.**
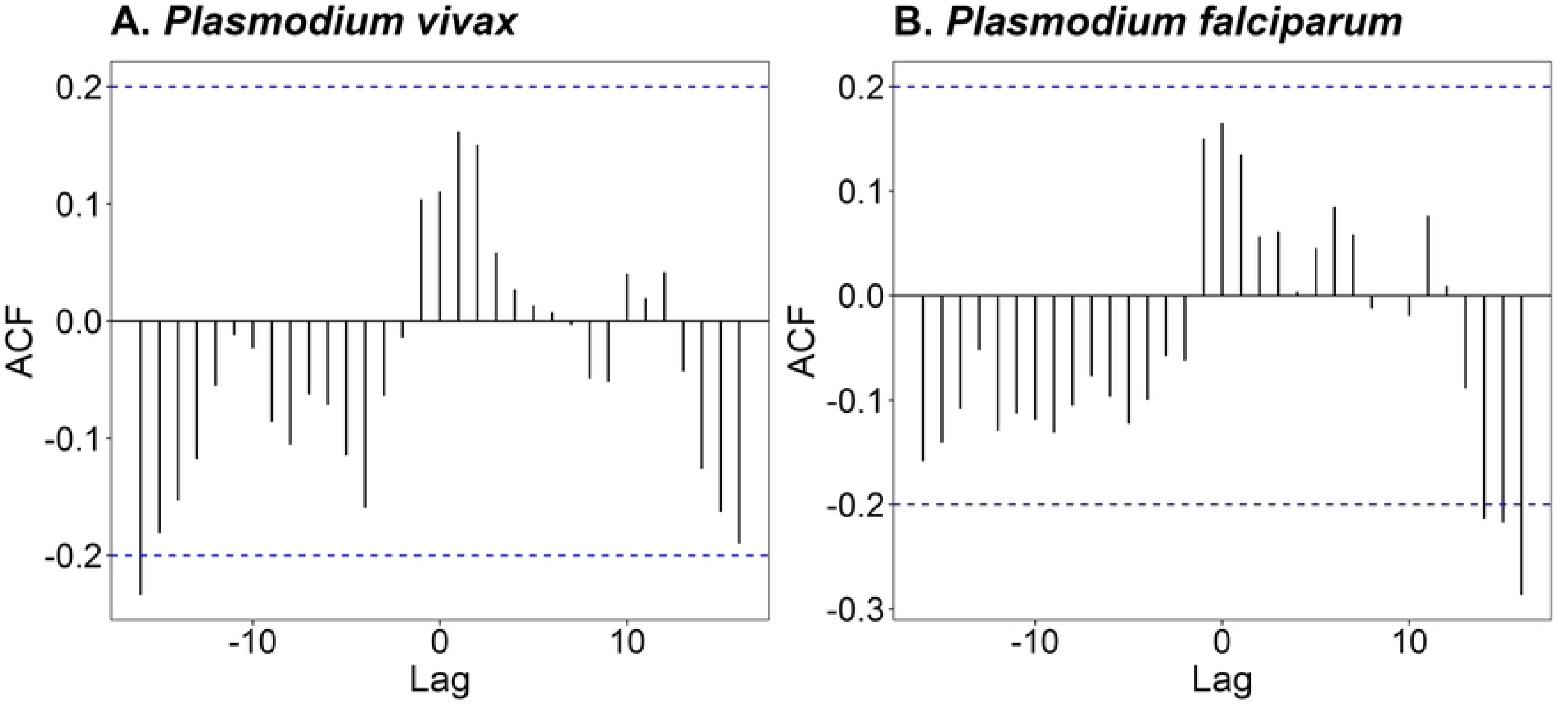
Anomalized rainfall and malaria incidence Rank Cross-Correlation. A. *Plasmodium vivax* incidence (PV). B. *Plasmodium falciparum* incidence (PF). ACF= Sample Cross-correlation Function.

**Fig 12.**
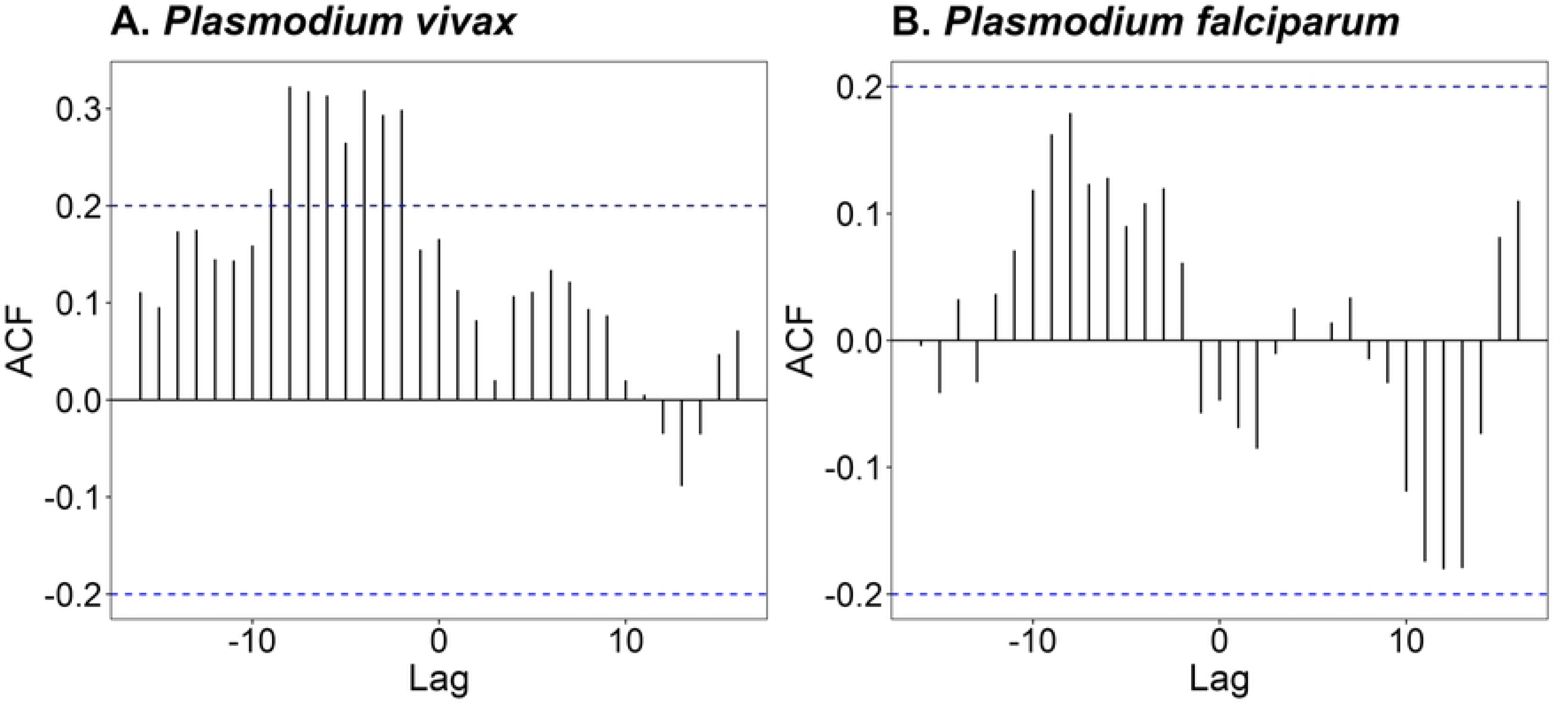
Anomalized temperature and malaria incidence Rank Cross-Correlation. A. *Plasmodium vivax* incidence. B. *Plasmodium falciparum* incidence. ACF= Sample Cross-correlation Function.

**Fig 13.**
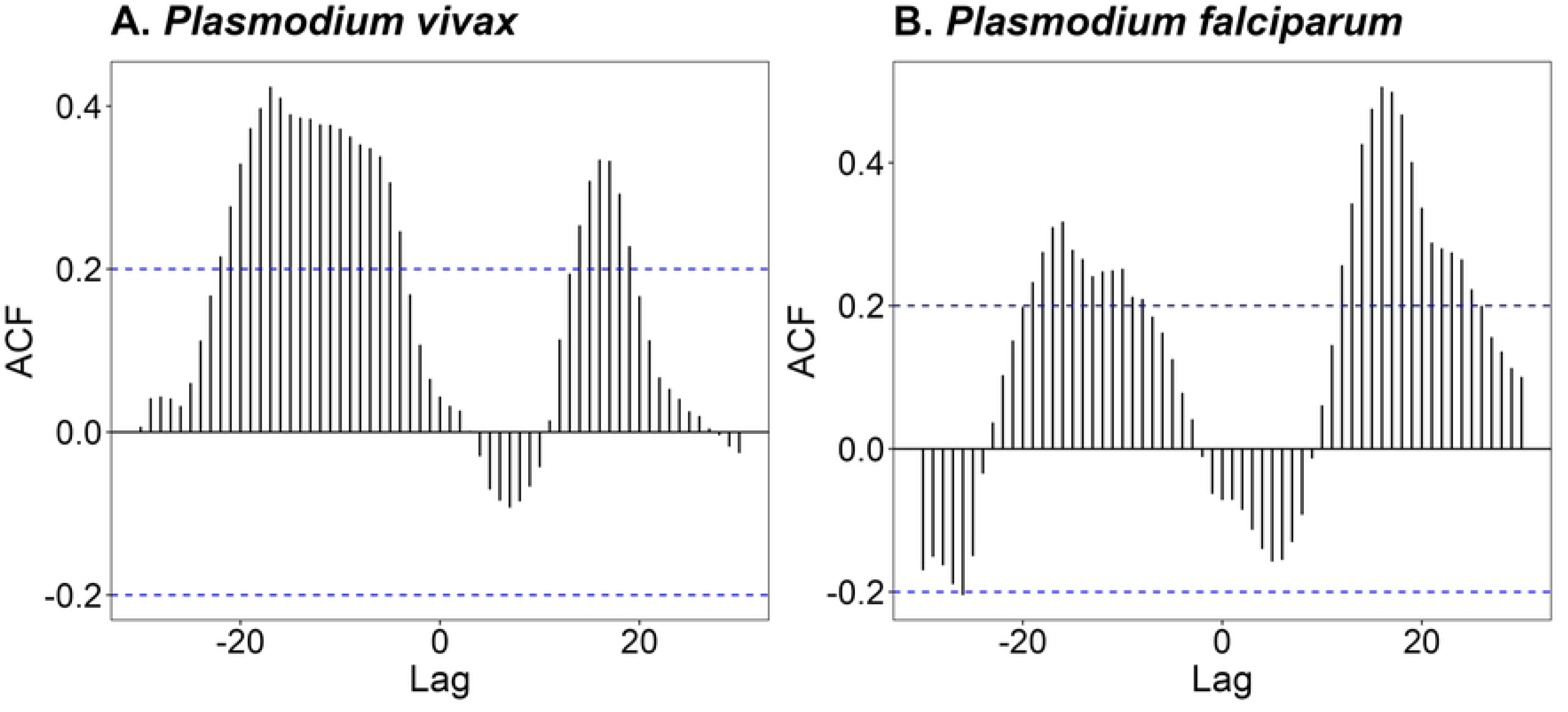
Anomalized El Niño 3.4 (ENSO) and malaria incidence Rank Cross-Correlation. A. *Plasmodium vivax* incidence (PV). B. *Plasmodium falciparum* incidence (PF). ACF= Sample Cross-correlation Function.

**Fig 14.**
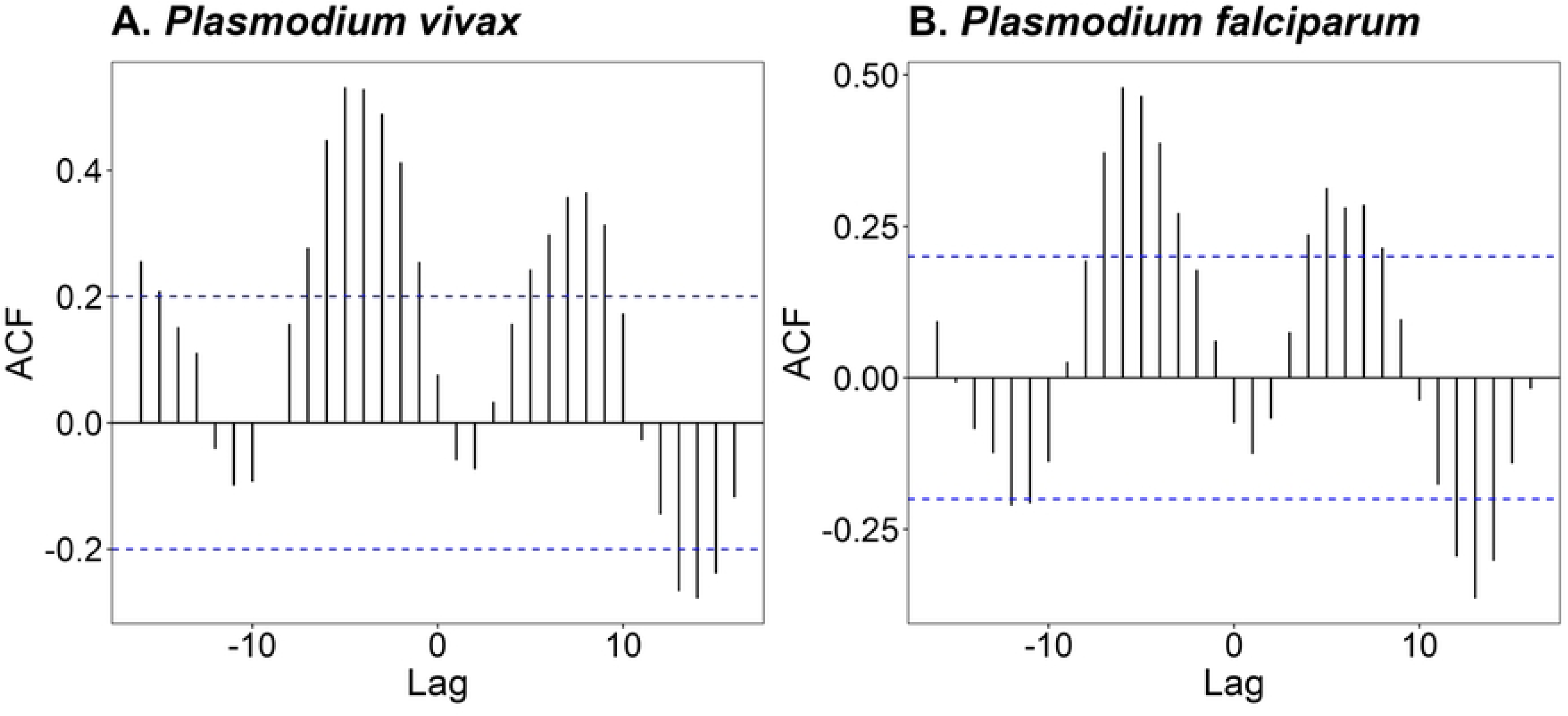
All *Anopheles* species counts and malaria incidence Rank Cross-Correlation. A. *Plasmodium vivax* incidence (PV). B. *Plasmodium falciparum* incidence (PF). ACF= Sample Cross-correlation Function.

**Table 3.** Selected lags for climate and mosquito count covariates to be included in the *Plasmodium vivax* and *Plasmodium falciparum* malaria incidence models for All *Anopheles* species and for *Anopheles darlingi*.

| Malaria<br>Model | Mosquito<br>Species | Rainfall | Temperature | ENSO | Mosquito<br>count |
| --- | --- | --- | --- | --- | --- |
| *P. vivax* | All species | 2, 4 | 8 | 17 | 4 |
| *P. falciparum* |  | 2, 5 | 8 | 16 | 5 |
| *P. vivax* | *An. darlingi* | 2, 4 | 8 | 17 | 4 |
| *P. falciparum* |  | 2, 5 | 8 | 16 | 6 |

Finally, we considered regressing the malaria incidence on past observations and past conditional means. The Sample Autocorrelation Function (ACF) and Partial Autocorrelation Function (PACF) plots of the malaria time series shown in Fig 15 was used to determine the lag of past observations that had the greatest impact on the malaria incidence cases at time *t*. Due to the highly significant partial autocorrelation at lag 1, we included past malaria observations at time *t* ― 1 only. Furthermore, after including the past observation, we also consider adding the conditional mean at *t* ― 12 to account for possible seasonal effects.

**Fig 15.**
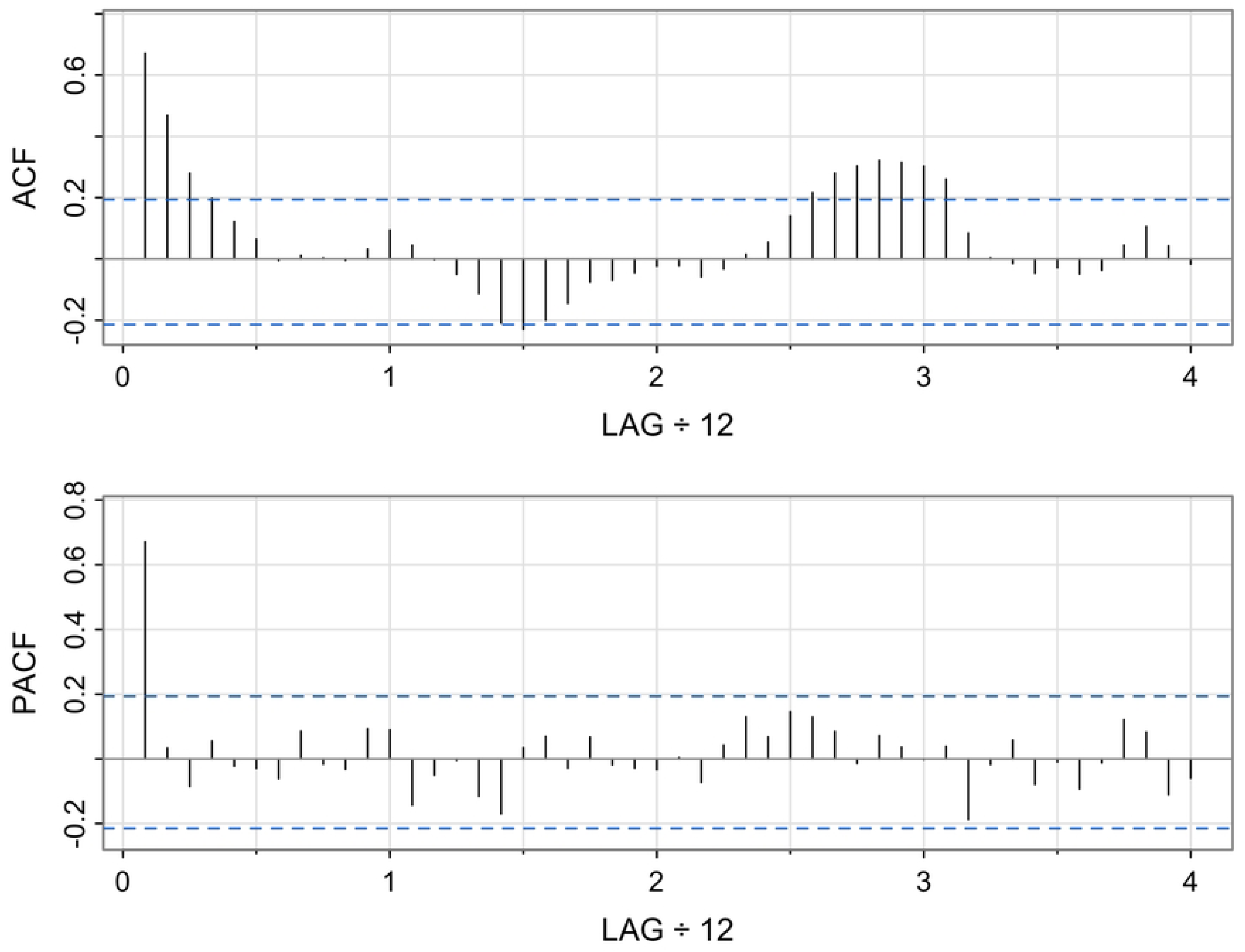
Sample Autocorrelation Function (ACF) and Partial Autocorrelation Function (PACF) for *Plasmodium vivax* incidence time series.

For all models, the logarithmic link function *g*(*λ*_*t*_) = *log*(*λ*_*t*_) was used. The identity function cannot fit models with negative covariates (such as the ENSO index) and cannot estimate effect parameters less than 0. Since some effects may be negative, the identity function is too restrictive in this case.

### Malaria Model Fitting and Prediction

The malaria model was fitted to the incidence of *P. vivax* and *P. falciparum*. To select the best Time Series Generalized Linear Model (TSGLM), we calculated different accuracy metrics for each unique combination of lagged and non-lagged covariates (rainfall, temperature, ENSO, and mosquito counts), an autoregressive term of malaria incidence at *t* ― 1, and the conditional mean malaria incidence at *t* ― 12. This term considers any seasonal dependence that might be present in the data. The conditional mean was only considered if the autoregressive term was already in the model. Mosquito counts for All *Anopheles* species imputed with the GB method were proposed as potential covariates. Additional results were also produced using *An. darlingi* mosquito counts as a predictor. We also included an additional trend shift component to account for the observed trend increase of malaria incidence from year 2015 onward.

We focus on selecting the model that gives the best prediction accuracy when compared to observations. For this purpose, we used an 80-20 training-testing sample split. The training set was used to fit models according to the above algorithm, and each model was used for prediction in the testing set. Using these predictions, we calculated a variety of error metrics: the root mean square error (RMSE), the mean absolute error (MAE), and the mean absolute percentage error (MAPE). Lower error values are preferred. The best models, as indicated by the lowest values of RMSE, MAE, and MAPE, are presented in Table 4.

**Table 4.** Selected covariates for the *Plasmodium vivax* and *Plasmodium falciparum* malaria incidence models that minimize the prediction error in a testing set using different accuracy measures.

| *Plasmodium vivax* |  |  |  | *Plasmodium falciparum* |  |  |  |
| --- | --- | --- | --- | --- | --- | --- | --- |
| Covariates | RMSE | MAE | MAPE | Covariates | RMSE | MAE | MAPE |
| Rainfall | X<sup>a</sup> | X | X<sup>a</sup> | Rainfall | X<sup>a</sup> | X<sup>a</sup> | X<sup>a</sup> |
| Rainfall (lag 2) |  | X | X | Rainfall (lag 2) | X | X<sup>a</sup> | X<sup>a</sup> |
| Rainfall (lag 4) | X | X | X | Rainfall (lag 5) |  |  |  |
| Temperature | X | X | X | Temperature |  | X | X |
| Temp (lag 8) |  |  |  | Temp (lag 8) | X |  |  |
| ENSO | X | X | X | ENSO | X<sup>a</sup> | X<sup>a</sup> | X<sup>a</sup> |
| ENSO (lag 17) |  |  |  | ENSO (lag 17) | X<sup>a</sup> | X<sup>a</sup> | X<sup>a</sup> |
| Mosquito Count<sup>b</sup> | X | X |  | Mosquito Count<sup>b</sup> |  |  |  |
| Mosq Count (lag 4) | X | X | X | Mosq Count (lag 4) |  |  |  |
| Past Obs<sup>c</sup> (lag 1) | X<sup>a</sup> | X<sup>a</sup> | X<sup>a</sup> | Past Obs<sup>c</sup> (lag 1) | X<sup>a</sup> | X<sup>a</sup> | X<sup>a</sup> |
| Past Mean (lag 12) |  |  |  | Past Mean (lag 12) | X | X | X |
| Time Shift<sup>d</sup> | X<sup>a</sup> | X<sup>a</sup> | X<sup>a</sup> | Time Shift<sup>d</sup> | X | X | X |

RMSE= root mean square error; MAE= mean absolute error; MAPE= mean
absolute percentage error.
<sup>a</sup> Coefficient is significant at 5%
<sup>b</sup> Mosquito counts include All *Anopheles* species imputed using Gradient Boosting
<sup>c</sup> Past Observation
<sup>d</sup> Time Shift Post-2015

The general malaria incidence model for *P. vivax* or *P. falciparum* can be written as:

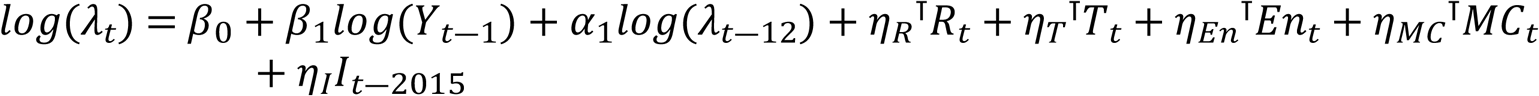

where *λ*_*t*_is the expected malaria incidence rate and *Y*_*t*_ is the observed incidence rate at time *t* [(cases/population) × 1,000]. *R*_*t*_, *T*_*t*_ and *En*_*t*_are the climatic variables (rainfall, temperature, ENSO respectively) vector with non-lagged and lagged values at time *t*. *MC*_*t*_is the mosquito counts vector with non-lagged and lagged values at time *t*. *I*_*t*―2015_ is an indicator variable which is zero if the year is lower than 2015 (constant trend), and it is proportional to time if time is greater than 2015. Model parameters *β*_0_ (intercept), *β*_1_ (*t* ― 1 observation), *α*_1_ (*t* ― 12 mean), and parameter vectors *η*_*R*_,*η*_*T*_,*η*_*En*_,*η*_*I*_are estimated by maximum likelihood using the R library *tscount* [43].

Selected covariates in Table 4 (marked with “X”) minimize the different prediction error metrics and may vary depending on the prediction criteria used. For the *P. vivax* malaria model, rainfall, lagged rainfall (lags 2 (MAE and MAPE) and 4), temperature, ENSO, mosquito counts (RMSE, MAE), lagged mosquito counts (lag 4) and the previous time-step expected malaria incidence are selected as the most important covariates in the sense of minimizing the predictive errors for the testing set, with a significant trend shift after the year 2015. Only rainfall, past month observations, and the trend shift component are significant at the 95% confidence level.

For the *P. falciparum* malaria incidence model, rainfall, lagged rainfall (lag 2), temperature (MAE and MAPE), lagged temperature (lag 8, RMSE), ENSO, lagged ENSO (lag 16), the previous time step malaria incidence observation, past year mean incidence and a linear trend shift are selected as the most important covariates that minimize the model predictive errors over the testing data set. Only rainfall and lagged rainfall, ENSO and lagged ENSO, and the previous month observation are statistically significant according to the 95% confidence intervals calculated using the normal approximation for the model parameter estimates. Predictors selected by minimizing the predictive errors are not necessarily going to be statistically significant according to the Wald test statistic (see Table 4). However, predictability performance was preferred over statistical significance in this case.

Fig 16 shows the model fitting and forecast results for the *P. vivax* malaria incidence model for All *Anopheles* species and *An. darlingi* counts as predictors. Model forecasts also include the 80% and 95% confidence intervals. Two model versions were fitted: the first one used imputed *An. darlingi* as a predictor; and the second one used All *Anopheles* species mosquito counts as a predictor.

**Fig 16.**
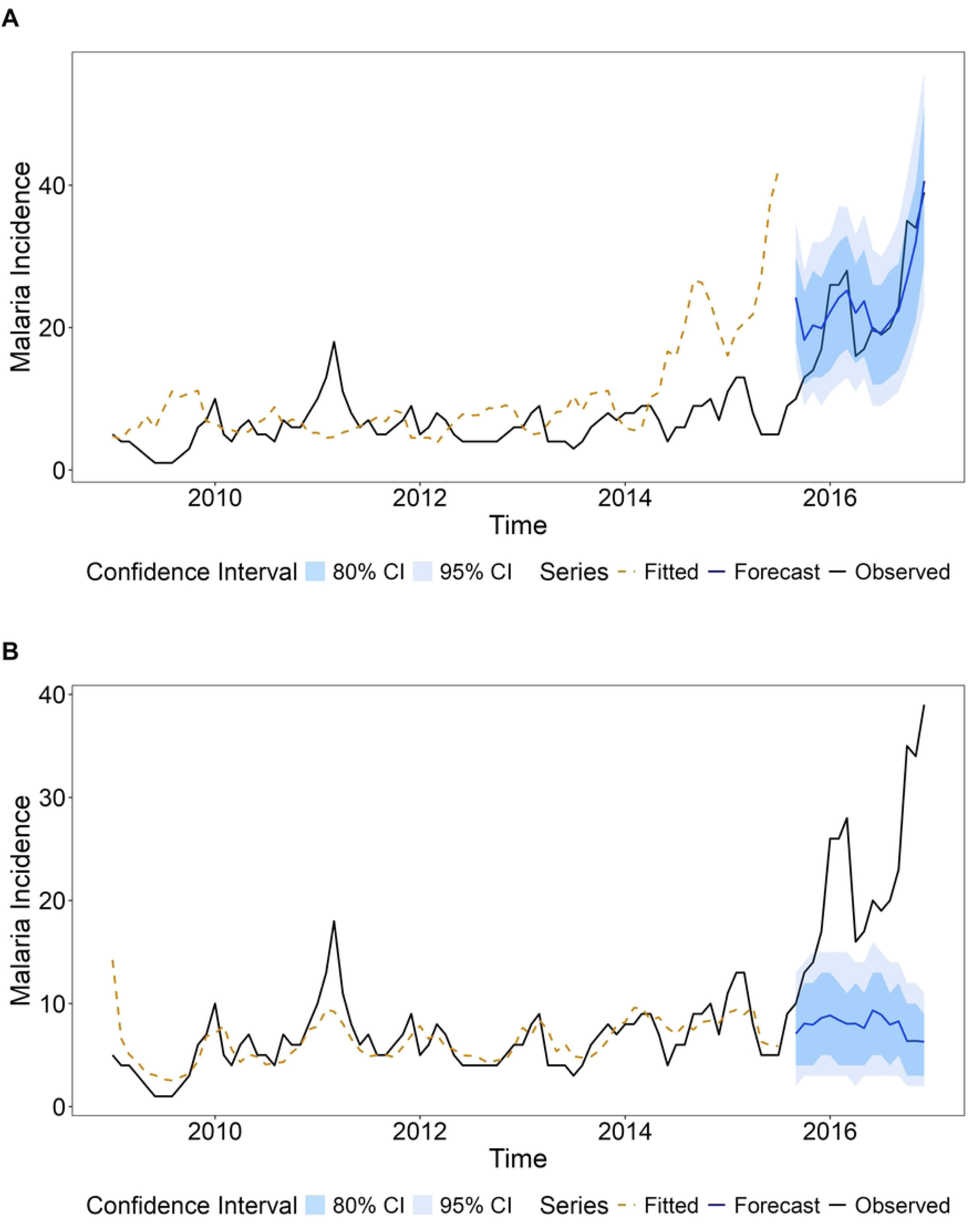
Model fit and predictions for *Plasmodium vivax* malaria incidence time series. A: All *Anopheles* species used as a covariate for mosquito counts. B: *Anopheles darlingi* used as a covariate for mosquito counts.

Fig 16 shows a better forecasting performance when using *All* mosquito species as a predictor (Fig 16A) in comparison when using *An. darlingi* (Fig 16B). In this case most of the observations in the testing set are included within the 80% confidence intervals with a clear upward trend from year 2015.

Fig 17 shows the model fitting and forecast results for the *P. falciparum* malaria incidence model. Based on the lowest prediction errors, the mosquito count variable was not included as a predictor (Table 4).

**Fig 17.**
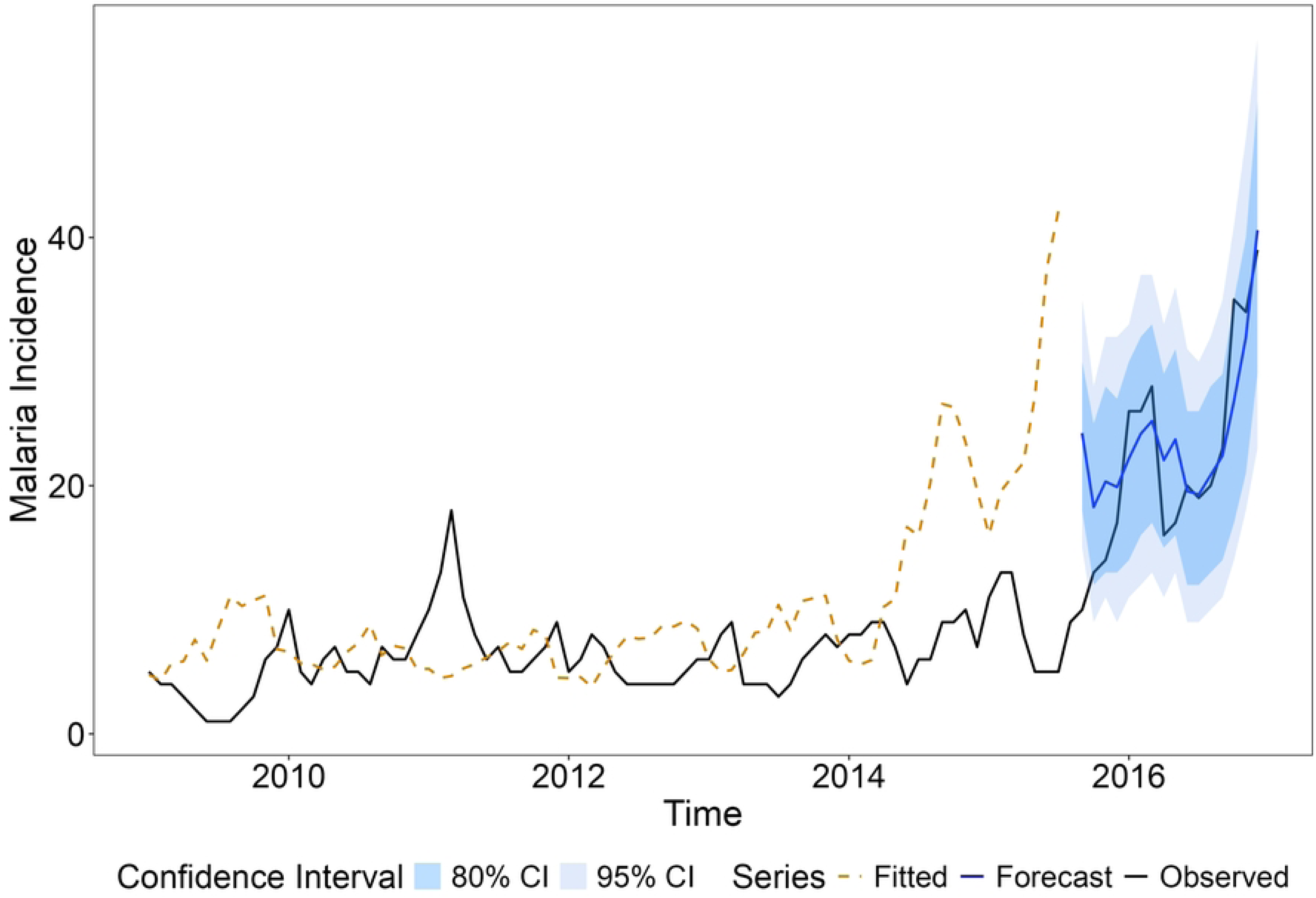
Model fit and predictions for *Plasmodium falciparum* malaria incidence time series.

According to Fig 17, data observations in the testing set are included within the 95% and 80% confidence intervals of the predicted values.

#### Malaria Model sensitivity to mosquito data imputation methods

An important question to be addressed is the sensitivity of the mosquito counts data imputation methods in the malaria model configuration and prediction accuracy. Although the Gradient Boosting approach resulted in the best imputation method performance according to the results presented in Table 2, mosquito counts imputed with all methods were tested as potential predictors, and variable selection was recorded in each case. We explored how predictability changes when using different mosquito imputed time series in the model selection process. Model configuration (or best predictors set) can also change when different imputed mosquito count time series are used in the model selection process.

Gradient Boosting (GB) imputed mosquito counts and lagged mosquito counts for All *Anopheles* species were not included as potential model covariates in the *P. falciparum* best predictive model; however, mosquito counts were selected as covariates in the *P. vivax* malaria model by all prediction accuracy criteria.

In Table 5, we compare the different accuracy measures in the 20% testing data set for the *P. vivax* and *P. falciparum* malaria models and the four different time series of imputed mosquito counts for All *Anopheles* species.

**Table 5.**
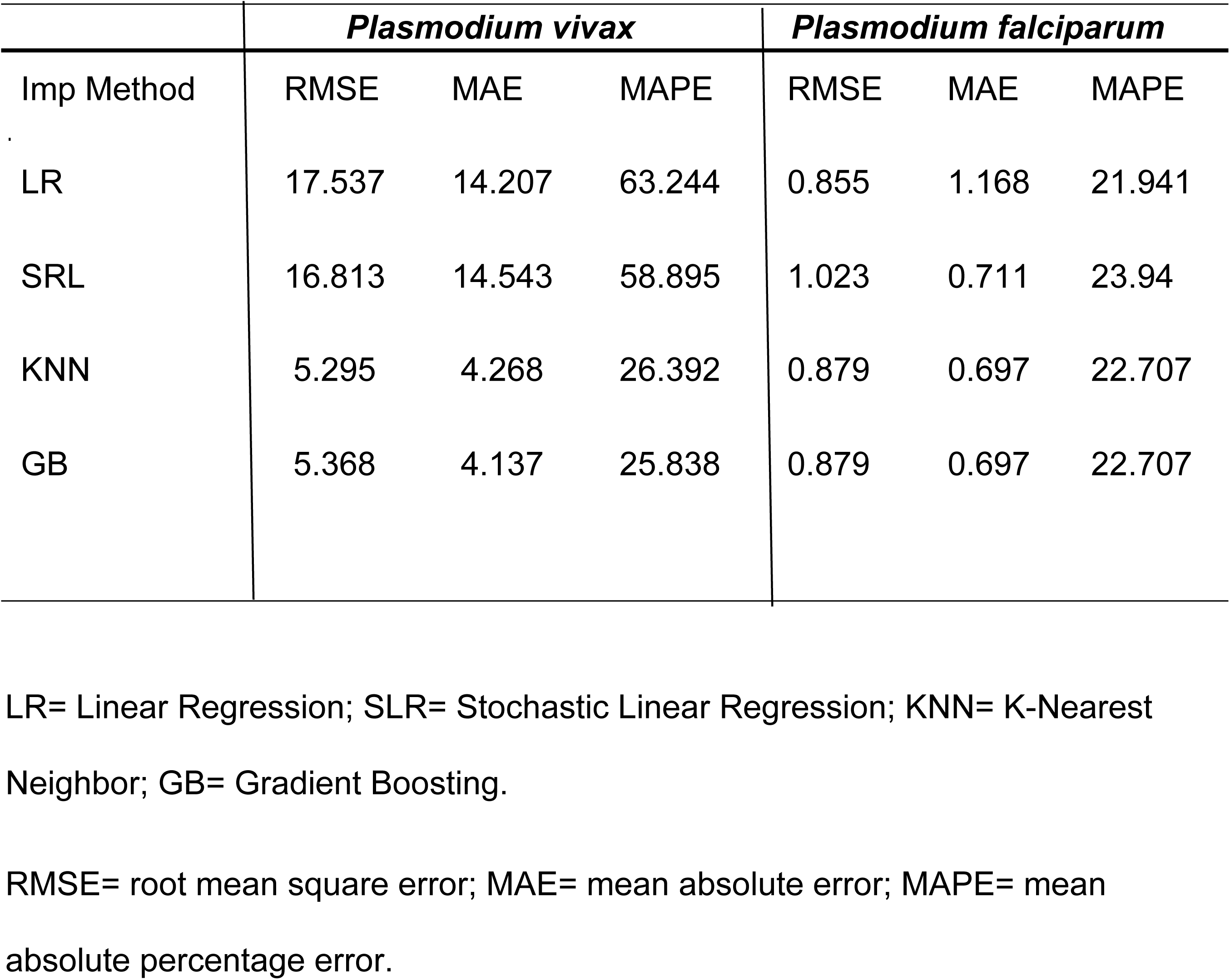
*Plasmodium vivax* and *Plasmodium falciparum* malaria models. Comparison of prediction errors for different imputation methods for mosquito counts time series for All *Anopheles* species.

|  | *Plasmodium vivax* |  |  | *Plasmodium falciparum* |  |  |
| --- | --- | --- | --- | --- | --- | --- |
| Imp Method | RMSE | MAE | MAPE | RMSE | MAE | MAPE |
| LR | 17.537 | 14.207 | 63.244 | 0.855 | 1.168 | 21.941 |
| SRL | 16.813 | 14.543 | 58.895 | 1.023 | 0.711 | 23.94 |
| KNN | 5.295 | 4.268 | 26.392 | 0.879 | 0.697 | 22.707 |
| GB | 5.368 | 4.137 | 25.838 | 0.879 | 0.697 | 22.707 |

LR= Linear Regression; SLR= Stochastic Linear Regression; KNN= K-Nearest Neighbor; GB= Gradient Boosting.
RMSE= root mean square error; MAE= mean absolute error; MAPE= mean absolute percentage error.

The results show that for the *P. vivax* malaria incidence model, the lowest prediction errors are obtained when imputed K-Nearest Neighbor (KNN) and Gradient Boosting (GB) All *Anopheles* species counts are used as covariates. Larger prediction errors are obtained when using imputed LR and SLR mosquito time series as covariates in the model, compared with using imputed GB and KNN mosquito time series.

The best predictive *P. falciparum* malaria incidence models exclude the covariate mosquito counts when K-Nearest Neighbor (KNN) and Gradient Boosting (GB) imputed time series are used as predictors. In this case, prediction errors for the malaria incidence model are the same for both time series. However, this result changed when Linear Regression (LR) or Stochastic Linear Regression (SLR) were used as imputation methods. All *Anopheles* species-imputed time series were included as potential covariates in the malaria incidence model in this case.

Prediction errors across each predictive measure (RMSE, MAE, MAPE) do not show high variability among different imputation methods.

Results from Table 5 indicate a greater sensitivity to the imputation methods for the *P. vivax* malaria incidence model than for the *P. falciparum* model.

## Discussion

The elimination of malaria requires a strong surveillance system. A key component is entomological surveillance which relies on monitoring anopheline populations mainly in terms of species composition, abundance, biting behavior, and insecticide resistance [48]. The collection of entomological data requires a huge investment in regular mosquito collections in various sentinel sites throughout endemic areas. Systematic data collection approaches as the one presented by Drakou et al [49] provides a good example of the implementation of mosquito network surveillance with adequate spatial coverage, while in our study implementing a comprehensive spatial surveillance effort was logistically unfeasible. In the study by Drakou et al [49], surveillance was limited to a single year. However, a multiyear analysis, such as the one we are presenting, is always valuable for understanding the impact of environmental factors on mosquito bionomics and to understand year-to-year variability.

In remote locations with difficult access, a way to contribute to providing relevant entomological data to the malaria program is through well-trained local leaders. This approach was attempted in an Amerindian village of the Caura River basin in southern Venezuela. Nevertheless, the political, economic, and social crisis [50,51] jeopardized the success of such an initiative, resulting in irregular interruptions of mosquito collections with the end result of large missing data, making it difficult to interpret the data and provide reliable information for decision-making for vector control.

To overcome this limitation, the application of data imputation methods arise as a potential solution. Machine learning approaches, specifically Boosting methods have been proposed by other authors when there is a significant percentage of missingness [4]. However, imputing missing data under the assumption that it is missing at random may not be entirely accurate, as intrinsic seasonal and operational factors may have influenced the sampling effort.

Given the large percentage of missing data in the mosquito abundance time series, the application of different imputation methods provides a good alternative when mosquito surveillance is highly fragmented, and continuous time series are not available. However, different imputation methods might capture different data features, and testing the feasibility of the imputed values still remains a challenging task. Four imputation methods were compared: Linear Regression, Stochastic Linear Regression, K-Nearest Neighbor, and Gradient Boosting.

Considering the limited data availability and rather short time series, the LOOCV approach was the best option to evaluate predictability among the different methods. Gradient Boosting and the Stochastic Linear Regression methods demonstrated better efficiency in terms of accuracy, reporting the lowest LOOCV errors (Table 2). In contrast, the Linear Regression approach was less efficient than the other methods, showing the highest LOOCV errors and a poorer representation of the seasonal cycle. Comparison among different imputation methods for mosquito data and any other time series data is a necessary first step in the evaluation of potential biases in the new data information provided [52]. If this new data is used as input to an impact model, conducting a sensitivity analysis of the imputation methods on the expected response would provide additional insights into the usefulness of the imputed data. In general, the seasonal cycle was well represented by all imputation methods, with marked seasonal peaks between August and September, two to three months after the beginning of the rainy season. Similar results were recorded in other malaria endemic areas of Venezuela [13,39], while in other regions the seasonal peaks are related to the transition period between rainy and dry season [38,53,54]. The large spike observed in 2010 (Fig 5), has been associated to a weak La Niña condition. A similar spike to the one observed in 2010 was estimated for 2016 when applying the GB imputation method to the *An. darlingi* time series, with a maximum comparable to the one recorded in 2010. However, mosquito measurements were only available through July 2016, so direct validation for this period is not possible. Furthermore, we cannot conclude that there is a direct association of this spike with a weak La Niña event, since the next event was reported during the Fall 2017 up to Spring 2018 [55]. It is important to point out that although there are abundant references worldwide relating malaria and other diseases to ENSO events, the available data on mosquito abundance and ENSO is scant. El Niño is associated with an increase in rainfall in the Colombia Pacific coast and East Africa, and La Niña with drought. A three-year study conducted in the Colombian Pacific coast reported no relation between ENSO and the abundance of *An. albimanus* and *An. darlingi* [56] while a one-year study in Tanzania showed a decline in the abundance of *An*. *arabiensis* and *An*. *funestus* associated with the drought due to La Niña 2016-2017 [57].

The relationship between climate and malaria is powerful, complex, and highly localized. Several studies clearly show that factors like temperature and rainfall, and the global climate patterns like El Niño that influence them, are critical drivers of malaria epidemics [17,58,59]). Previous studies in Venezuela have shown that El Niño is associated with drought, resulting in an increase in malaria cases the following year [21,22,23]). In the present study the significant lags for rainfall and ENSO were one month earlier for the *P. vivax* model than for the *P. falciparum* model (Table 3), which is reasonable considering that the sporogonic cycle of *P. falciparum* is larger than that of *P. vivax* [60,61]. Although variations in temperature have a great impact on the developmental time of the mosquitoes and the parasites [60,61,62], the lag of 8 months was the most significant for both models. When fitting the *P. vivax* and *P. falciparum* malaria incidence models, the main issue is the non-significance or exclusion of mosquito abundance as covariates for the *P. falciparum* malaria incidence model. These findings suggest that it is insufficient to treat mosquito abundance and diversity data collected from a single locality as representative for the entire municipality, because these ecological characteristics can differ significantly between localities within the same region [11,38,39,53].

However, the proposed malaria model is still useful for predictive purposes, since the model accounts for potential seasonal effects and significant changes in the incidence trend, lagged and non-lagged environmental factors, and the impact of the previous month’s malaria incidence. The better model performance observed when using All *Anopheles* species as a mosquito count predictor, compared with using *An. darlingi* (Fig 16) might be due to the unexpected spikes obtained in the imputed mosquito count time series of *An. darlingi* for the Boosting method (GB) (Fig 10D).

The sensitivity analysis using different imputation methods is a key component in the results presented in this work. These results are demonstrated through the process of evaluating both model accuracy and model structure, as these aspects are sensitive to the imputation method employed. Therefore, it is important to explore all available methods to ensure robust and reliable conclusions. When assessing the sensitivity of the different imputation methods on the malaria model predictive performance, the MAPE results shown in Table 5 confirms a reasonably good predictive performance of the *P. vivax* model when mosquito counts time series for All *Anopheles* species imputed with KNN and GB methods are used as predictors (20% <MAPE<30%), while values of MAPE are greater than 50% when mosquito count time series imputed with LR and SLR methods are used instead. Thresholds to evaluate model predictive performance according to the MAPE criteria have been discussed by Ma and Liu [63].These results might be an indication that these two imputation methods (LR and SLR) do not accurately represent the mosquito abundance variability for All *Anopheles* species, and as a consequence might fail to reproduce the association between mosquito abundance and malaria incidence. In the case of the *P. falciparum* model predictive performance according to MAPE is more homogeneous across the different imputation methods (Table 5). But in this case mosquito abundance does not yield a better model accuracy when the KNN and GB imputed mosquito counts are considered as predictors. Differences in model accuracy between *P. vivax* and *P. falciparum* are driven by the selection of the sets of predictors that minimize prediction errors. Mosquito counts imputed with the regression models for All *Anopheles* species preserve the seasonality in the mosquito abundance cycle but fail to represent important year-to-year variability fluctuations exhibiting a higher degree of smoothing. However, when used as predictors they can explain a good percentage of the malaria time series variability in the *P. falciparum* model. This is not the case for the GB and KNN imputed mosquito time series, for which the more complex fluctuations do not contribute to explain the observed variance in the malaria time series model. We can speculate that data limitations may be an important factor underlying these results.

Despite data limitations, this work provides a template for integrating multiyear entomological observations, climate information, and robust imputation techniques to improve malaria prediction systems in remote settings. The approaches developed here can support more timely and geographically informed decision-making, enabling public health authorities to anticipate high-risk periods, optimize vector control interventions, and better allocate scarce resources. Continued investment in local capacity for mosquito monitoring—combined with modern data-analytic tools—can strengthen the foundations of malaria surveillance and contribute to more effective control strategies in the Amazon and similar hard-to-reach regions.

Finally, a key entomological parameter for the implementation of a particular intervention for control is to determine the vector biting hourly pattern. This pattern is species specific; nevertheless, *An. darlingi* exhibits different patterns along its range of distribution, even among locations within a few kilometers apart [11,64,65,66]. During the present study, the mosquito abundance at different hour intervals was not significant, contrasting with previous studies in Sucre municipality along the Caura River where *An. darlingi* showed a biting peak around midnight in Las Majadas [64] whereas in Jabillal, there were two distinctive peaks at sunset and sunrise [11]). The lack of significance in the hour intervals might be due to the low abundance of mosquitoes collected throughout the study. Similar results were reported for *An. calderoni* in Colombia [67].

## Conclusion

This study demonstrates that machine learning–based imputation methods offer a powerful tool for reconstructing highly incomplete mosquito abundance time series in remote, resource-limited malaria-endemic regions. Incorporating the imputed mosquito series and climate predictors into generalized time-series models of malaria incidence revealed important differences between *P. vivax* and *P. falciparum*. Models for *P. vivax* responded strongly to mosquito abundance, rainfall, temperature, ENSO anomalies, and recent malaria history, with Gradient Boosting and KNN imputations yielding the most accurate forecasts. In contrast, *P. falciparum* incidence was largely insensitive to mosquito abundance, likely reflecting mismatches between the geographic scale of entomological and epidemiological datasets, lower case counts, or intrinsic differences in transmission. Climate drivers—particularly rainfall and ENSO—retained predictive power for both parasites, underscoring their central role in shaping transmission potential.

## Acknowledgment

We are most grateful to Simón Caura, the local leader, Hernán Guzmán, Yarys Estrada, Victor Sánchez, Jorge Moreno and Jesús González, who participated in field activities. To the people of Boca de Nichare, for their support and friendship. To Ana María Ibañez and Horacio Vargas for logistic support and friendship. Mariapia Bevilacqua, Lya Cárdenas, and Zenaida Muria (ACOANA) and Dirección de Salud Ambiental (MPPSalud) for logistic support.

## Supporting information

**S1 Fig. El Niño 3.4 index for the period 2009 – 2016.**

**S2 Fig. Root mean square prediction error (RMSE) after 1,000 iterations of the random sampling approach [leave-one-out cross-validation (LOOCV)] for each Anopheles species, showcasing the impact of lagging/no-lagging of climatic variables using an approach with linear regression imputation.** All= all Anopheles species collected; darl.1= Anopheles darlingi; goel.6= Anopheles goeldii; osw.5= Anopheles oswaldoi; other= other Anopheles species collected.

**S3 Fig. Root mean square prediction error (RMSE) for each species using a leave-one-out cross-validation (LOOCV) approach with stochastic linear regression imputation.** All= all *Anopheles* species collected; darl.1= *Anopheles darlingi*; goel.6= *Anopheles goeldii*; osw.5= *Anopheles oswaldoi*; other= other *Anopheles* species collected.

**S4 Fig. Estimated total number of All *Anopheles* species abundance (ETNM) time series Gradient Boosting (GB) imputation method.**

